# Endosome dysfunction leads to gain-of-function TLR7 and human lupus

**DOI:** 10.1101/2023.04.03.535356

**Authors:** Harshita Mishra, Claire Schlack-Leigers, Ee Lyn Lim, Oliver Thieck, Thomas Magg, Johannes Raedler, Christine Wolf, Christoph Klein, Helge Ewers, Min Ae Lee-Kirsch, David Meierhofer, Fabian Hauck, Olivia Majer

## Abstract

Hyperactive Toll-like receptor (TLR) 7 signaling has long been appreciated as a driver of autoimmune disease by breaking tolerance to self-nucleic acids in mouse models^1–5^. Recently, mutations in TLR7 or its associated regulator UNC93B1^6, 7^, were identified as monogenic causes of human lupus; the unifying feature of these mutations being TLR7 gain-of-function. TLR7 is an intracellular transmembrane receptor, sensing RNA breakdown products within late endosomes^8, 9^. Hence, its function depends on intricate transport mechanisms and membrane interactions within the endomembrane network. Whether perturbations of any of these endosome-related processes can give rise to TLR7 gain-of-function and facilitate self-reactivity has not been investigated. Here, we show that a dysregulated endosomal compartment leads to unrestricted TLR7 signaling and human lupus. The late endosomal BLOC-1-related protein complex (BORC) together with the small Arf1-like GTPase Arl8b controls TLR7 protein levels, and a direct interaction between Arl8b and Unc93b1 is required to regulate TLR7 turnover. We identified an amino acid insertion in UNC93B1 in a patient with childhood-onset lupus, which reduces the interaction with the BORC-Arl8b complex and leads to endosomal TLR7 accumulation. Therefore, a failure to control the proper progression of TLR7 through its endocytic life cycle is sufficient to break immunological tolerance to nucleic acids in humans. Our results highlight the importance of an intact endomembrane system to prevent autoimmune disease. As the cellular mechanisms restricting TLR7 signaling can be manifold, identifying and stratifying lupus patients based on a TLR7-driven pathogenesis could be a viable strategy towards a targeted therapy.

## Main

Mechanistic studies on TLR7 gain-of-function have primarily focused on hard-wired mutations in either TLR7, or its associated trafficking factor Unc93b1. Known sources for a hyperactive TLR7 include *Tlr7* gene duplications^1,4,5^, mutations increasing ligand affinity^6^ or Unc93b1 mutations releasing the brake on TLR7 signaling^2,3,7^. However, regulatory mechanisms of TLR7 activity involving the receptor-containing organelle have not been described. TLR7 signals from late endosomes/lysosomes, the main degradative organelle in eukaryotic cells that also serves as a central coordinator and signaling hub of the cell^10^. Accessory and transmembrane proteins assemble in functional complexes at the endosomal membrane to orchestrate the response of key cellular processes to environmental cues. One of these complexes is BORC (BLOC1-related complex), a critical regulator of lysosome biology, that controls lysosome positioning within the cytoplasm, as well as vesicle fusion of late endosomes and autophagosomes with lysosomes^11^. BORC is composed of eight conserved subunits that assemble on the cytosolic surface of the endosome^12^. The complex recruits the small Arf1-like GTPase Arl8b that serves as an adaptor for connecting BORC to different downstream protein machineries to mediate vesicle trafficking and fusion. Here we describe how defective BORC-mediated degradative sorting of TLR7 can lead to unrestrained signaling and human lupus.

## Results

### The BORC-Arl8b complex restricts TLR7 signaling

The BORC complex has been identified and best studied in Hela cells. There, loss of BORC-Arl8b abrogates anterograde lysosomal transport and results in the clustering of late endosomes/lysosomes in the perinuclear region^12^, whereas overexpression relocates late endosomes into the cell periphery^13^. We first confirmed these results by either knocking down BORC subunit 5 (Borcs5) or overexpressing Arl8b (Fig. 1a). To examine how these endosome manipulations would affect endosomal TLR signaling, we co-expressed and stimulated TLR7 in these cells. BORC-depleted Hela cells showed enhanced TLR7 signaling, as measured by increased induction of endogenous *IL-8* in response to R848 stimulation (Fig. 1b). In contrast, overexpression of Arl8b suppressed TLR7 signaling, without affecting protein levels (Fig. 1c). As gain-of-function TLR7 is a known driver of autoimmune disease^6^ and juxtanuclear endosome clustering has been described in inflammatory pathologies, including neurodegenerative and lysosomal storage disorders^14–16^, we further investigated the BORC-mediated restriction of TLR7 signaling.

**Fig. 1:**
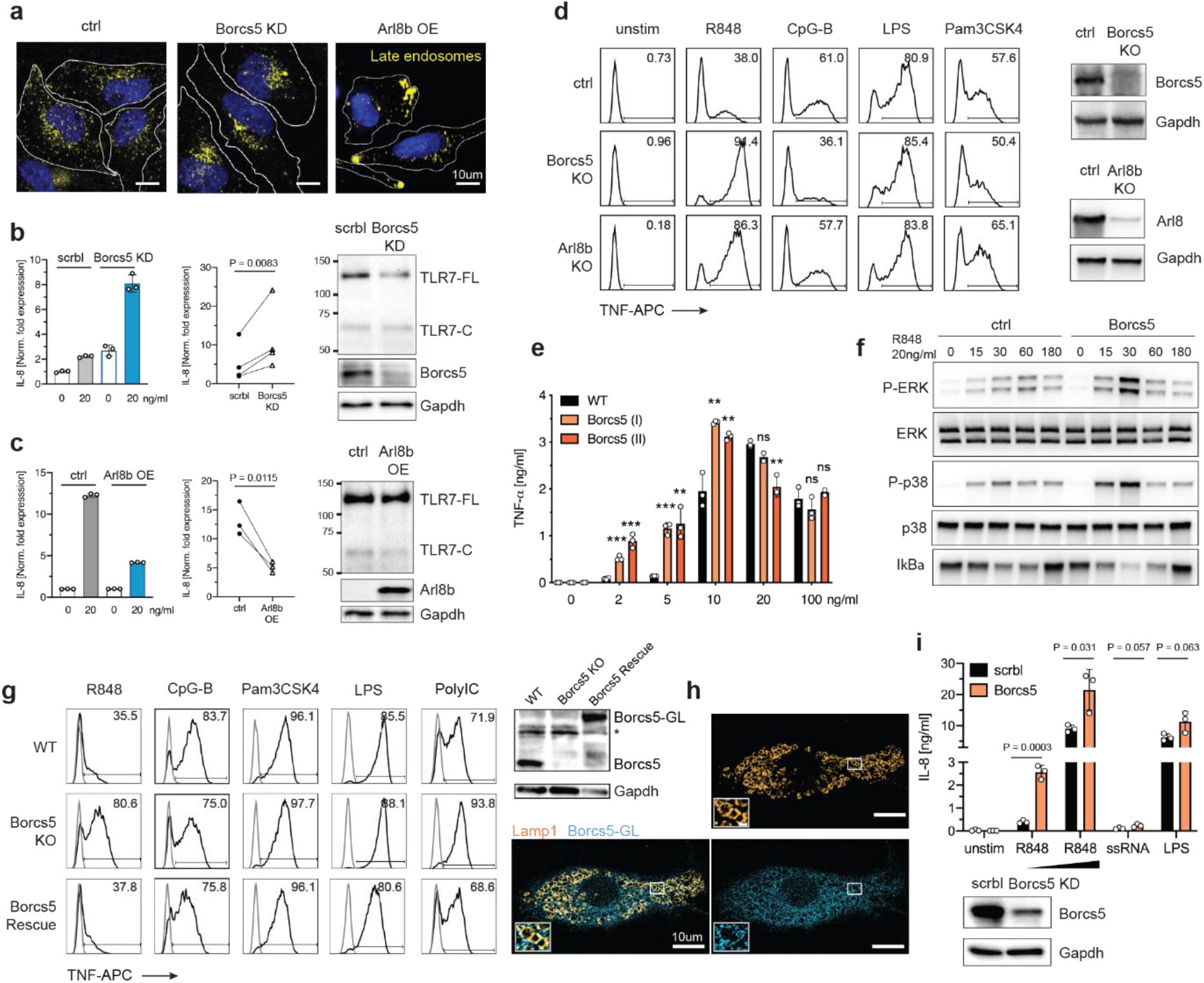
BORC-Arl8b restricts TLR7 signaling in human and mouse macrophages. **a**, Endosomes of Hela cells were loaded with AlexaFluor 488-dextran and transfected with either siRNA against Borcs5 or an Arl8b expression plasmid. Representative images showing the repositioning of late endosomes. **b**, Hela cells were co-transfected with TLR7-HA and siRNA against Borcs5. *IL8* gene induction was quantified by qPCR after 2h stimulation with R848. Bar graph shows a representative experiment. Dot plot shows pooled data of N=4; ratio paired t test, two-tailed. Western blots show TLR7-HA expression. C: cleaved, FL: full-length. **c**, Hela cells were co-transfected with TLR7-HA and Arl8b and treated as in **b**. Graphs displayed and analyzed as in B. **d**, Intracellular TNF staining of indicated Hoxb8 macrophage lines with R848 (10 ng/ml), CpG-B (300 nM), LPS (25ng/ml) or Pam3CSK4 (50ng/ml). Representative data of N=5. Western blots show KO validations. **e**, TNF ELISA of two Borcs5 KO Hoxb8 macrophages lines targeted with two separate guides after 8h stimulation with increasing concentrations of R848. Representative data (n=3) of N=5; unpaired t test, two-tailed. **f**, Immunoblot of phosphorylated P-p38, P-ERK and IκBα of Hoxb8 macrophages stimulated with R848 for the indicated times. Representative data of N=3. **g**, Intracellular TNF staining of indicated Raw246.7 macrophage lines with R848 (10 ng/ml), CpG-B (300 nM), Pam3CSK4 (50ng/ml), LPS (25ng/ml), or PolyIC (500ng/ml). Western blots show KO validation and rescue (* unspecific band). Representative data of N=3. **h**, Representative image of Raw246.7 expressing Borcs5-GreenLantern (GL) stained for GFP and Lamp1. **i**, IL8 ELISA of Borcs5 knock-down THP1 macrophages after overnight stimulation with increasing doses of R848 (2.5 and 5ng/ml), ssRNA (5µg/ml), and LPS (50ng/ml). Representative data (n=3) of N=2; unpaired t test, two-tailed. Western blot shows knock-down efficiency. \*\**P* < 0.01 and \*\*\**P* < 0.001.

To test if BORC would limit TLR7 signaling in immune cells, we deleted Borcs5 and Arl8b in mouse macrophages derived from Hoxb8-immortalized myeloid progenitor cells (hereafter referred to as ‘Hoxb8 macrophages’). Lentiviral CRISPR/Cas9-mediated knockout of Borcs5 or Arl8b greatly enhanced TLR7 signaling, as evidenced by higher production of TNF (Fig. 1d). The hyperresponsiveness was specific to TLR7, and did not apply to TLR9. As expected, surface TLR4 and TLR2 signaling were not affected by the endosome manipulation. The TLR7 hyperresponsiveness was dose-dependent and best demonstrated at suboptimal ligand concentrations (Fig. 1e). It was validated in two independent Borcs5 KO lines targeted with different guide RNAs (Fig. 1e) and was also observed when knocking out other BORC subunits, including Borcs6 and Borcs7 (Fig. S1a). The enhanced TNF response could be reverted to WT levels by reintroducing Borcs5 or Arl8b into the respective KO lines (Fig. S1b).

TLR7 was also more active in response to its physiological ligand ssRNA (Fig. S1c). BORC deletion led to increased phosphorylation of the MAPKs p38 and ERK, and stronger degradation of I*κ*B*α* downstream of TLR7 receptor activation (Fig. 1f), which was not observed when stimulating TLR9 (Fig. S1d). As all activated pathways returned to baseline with similar kinetics, negative feedback mechanisms were excluded. Enhanced TLR7 signaling was not a result of enriched levels of signaling molecules within the pathway, as proteomic analysis revealed normal levels of these proteins in BORC-deleted Hoxb8 macrophages (Fig. S1e).

To verify the phenotype in an independent macrophage system, we knocked out Borcs5 in murine Raw264.7 macrophages through Cas9-RNP delivery and obtained two independent clones. In both BORC-deleted clones, TLR7 signaling was enhanced, whereas TLR9 signaling was slightly reduced (Fig. 1g and Fig. S1f). Again, surface TLR2 and TLR4 were unaffected. The TLR7 hyperresponse was not restricted to the NFκB signaling pathway (TNF), but also extended to the type I interferon signaling branch (*Ifnb*) (Fig. S1g). We extended our analysis to TLR3, another endosomal RNA sensor, and also found this receptor to be more active in response to Poly-IC stimulation, although to a lesser degree than TLR7. All observed TLR signaling phenotypes could be rescued by reintroducing Borcs5 (Fig. 1g). Both mouse Borcs5 isoforms were able to rescue the TLR7 hyperresponse (Fig. S1h and 1i), even though only isoform 2 can be myristoylated, a modification believed to mediate lysosomal membrane attachment^12^. Imaging confirmed that both isoforms localized to late endosomes (Fig. 1h and Fig. S1j), suggesting that myristoylation is not the only way to anchor Borcs5 to endosomal membranes.

Finally, we verified our findings in human THP-1 macrophages. siRNA-mediated knockdown of Borcs5 in PMA-differentiated THP-1 macrophages corroborated the increased response to R848 and ssRNA40 seen in mouse macrophages, without affecting LPS signaling. (Fig. 1i).

These data show that the late endosomal BORC-Arl8b complex restricts TLR7 signaling in mouse and human macrophages.

### BORC deficiency has no effect on endosome positioning in macrophages

As BORC deficiency induces striking endosome clustering in epithelial cells, we next examined endosome positioning in macrophages. Contrary to Hela cells, endosomes in Arl8b- or Borcs5-deficient Hoxb8 macrophages did not collapse into the perinuclear region (Fig. 2a). To assess potential small differences in localization, we quantified the radial distribution of Lamp1^+^ late endosomes/lysosomes. There were no significant differences in the Lamp1 distribution in Borcs5, Borcs7 or Arl8b KO Hoxb8 macrophages compared to control cells. (Fig. 2b and Fig. S2a). We also assessed endosomal pH and proteolysis, as these are prerequisites for cleavage of endosomal TLRs to generate functional receptors^17^. The proteolytic capacity of endosomes in BORC-deficient Hoxb8 macrophages did not differ from control cells, as shown by comparable degradation of internalized DQ-BSA, a fluid-phase lysosomal proteolysis sensor (Fig. 2c). Equal endocytic uptake of cells was verified with fluorescent dextran (Fig. 2d). Also, luminal pH did not differ, as measured by the mean fluorescent intensity of internalized pHrhodo, a lysosomal pH reporter (Fig. S2b). To rule out differences in the overall number of late endosomes, we compared the amount of Lamp1 in KO and control Hoxb8 macrophages (Fig. S2c).

**Fig. 2:**
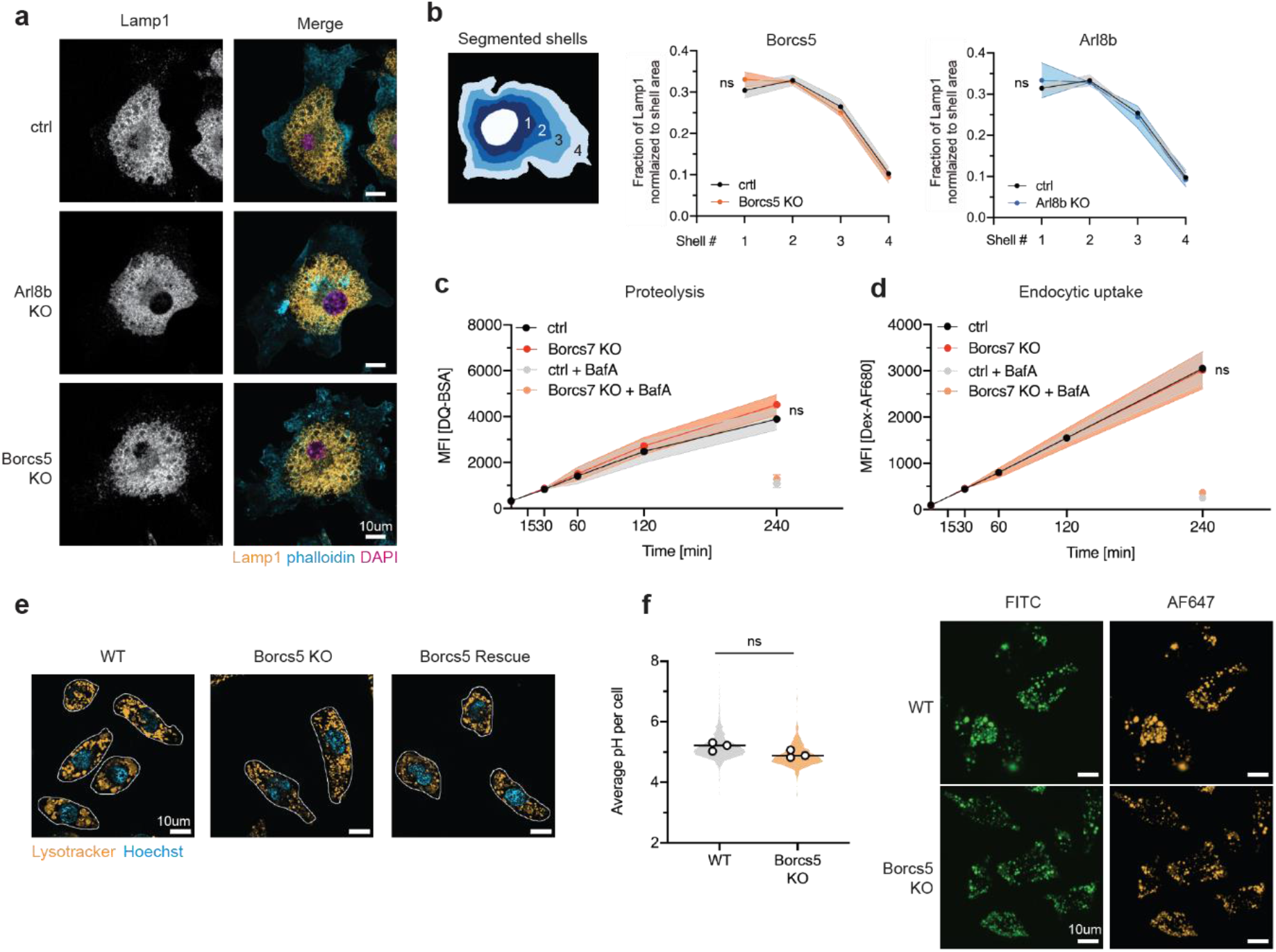
BORC-deficiency does not alter endosome positioning, luminal pH or proteolysis in macrophages. **a**, Indicated Hoxb8 macrophage lines were fixed and stained with anti-Lamp1, phalloidin, and DAPI to show endosome positioning. **b**, Quantification of endosome radial distribution across segmented shells. Graphs show the fraction of late endosomes normalized per shell area. Pooled data from N=3 (Borcs5) and N=2 (Arl8b), shaded areas indicate the SD. (Total number of cells analyzed: WT: 241, Borcs5 KO: 243, Arl8b KO: 163); two-way ANOVA comparing variation between genotypes. **c**, Quantification of the proteolytic capacity of endosomes by loading macrophages with the proteolysis reporter DQ-BSA Red for the indicated amounts of time. Bafilomycin A treatment served as negative control. Pooled data of N=3, shaded areas indicates the SD; two-way ANOVA comparing variation between genotypes. **d**, Quantification of endocytic uptake by loading macrophages with AlexaFluor680-Dextran for the indicated amounts of time. Data repeats and analysis as in C. **e**, Live cell images of indicated Raw246.7 macrophage lines fed with lysotracker, CellMask and Hoechst to show endosome positioning. **f**, Lysosomal pH of WT and Borcs5 KO Raw246.7 macrophages was determined by ratiometric fluorescence microscopy. Pooled data from N=3. Left: Dots represent means of independent experiments, shaded violin shows the variation of individual cells across all experiments; paired t test, two-tailed. (n = 982 WT & 884 Borcs5 KO cells). Right: Representative confocal images with identical acquisition and display settings.

We extended our analysis to Borcs5-deficient Raw264.7 macrophages and confirmed a similar endosome distribution with live-cell microscopy (Fig. 2e). To quantitatively assess the pH of single endosomes we performed ratiometric fluorescence microscopy. The luminal pH of individual endosomes did not significantly differ between WT and KO cells (Fig. 2f and Fig. S2d). As macrophages were plated on fibronectin-coated coverslips for this assay to improve cell spreading, we confirmed the hyperactive TLR7 response in Borcs5 KO cells on that coating (Fig. S2e).

Lastly, we assessed autophagy in BORC-deficient macrophages. Autophagy has previously been linked to TLR regulation^18^; and BORC promotes autophagy through facilitating the final fusion between autophagosome and lysosome^19^. We neither detected differences in the basal level of LC3B-II nor in the autophagic flux in Borcs5 KO Raw264.7 macrophages (Fig. S2f); excluding a role for autophagy in the BORC-mediated restriction of TLR7 signaling.

### BORC controls TLR7 turnover

As BORC deficiency did not visibly change endosome positioning, pH, proteolysis, or autophagy in macrophages, we compared the protein composition of endosomes from WT and Borcs5 KO Raw264.7 macrophages to identify other endosome-related processes that could functionally explain the TLR7 gain-of-function. First, we magnetically purified phagosomes from macrophages that had been fed iron beads and analyzed the phagosome proteome by mass spectrometry (Fig. 3a). Gene Set Enrichment Analysis revealed a broad reduction of lysosome-related proteins in the phagosomes of Borcs5 KO macrophages, including cathepsins, DNASE2, NPC2, and degradative enzymes of complex lipids and carbohydrates, indicating a major defect in this compartment (Fig. 3b and Fig. S3a). In agreement with the enhanced signaling response, TLR7 and TLR3 were enriched in phagosomes from KO cells. Surprisingly, there was also an enrichment of early endosomal markers and effector proteins involved in the Rab5-to-Rab7 conversion^20^, suggesting a defect in endosome maturation (Fig. 3b). Early endosomal markers were not expected in these preparations, as beads were given enough time to traverse all the way into lysosomes. We independently validated these findings via western blot and showed an accumulation of cleaved TLR7 and early endosomal markers, EEA1 and Rab5, in phagosomes from Borcs5 KO macrophages, which returned back to WT levels in the rescue line (Fig. 3c). A similar accumulation of TLR7 and EEA1 was observed in phagosomes purified from Borcs7-deficient Hoxb8 macrophages (Fig. S3b). The co-occurence of both late and early endosomal markers in phagosome preparations from KO cells suggested either the existence of an unusual hybrid compartment, decorated by both markers, or a delay in phagocytosis with beads retained in early endosomes at the time of harvest. To distinguish between these two possibilities, we imaged individual bead-containing phagosomes in BORC KO cells after 3h of synchronized uptake. Beads taken up by WT cells were exclusively present in Lamp1^+^ late endosomes/lysosomes. In Borcs5 KO macrophages a sizable portion of beads were found within hybrid compartments, marked by both EEA1 and Lamp1 (Fig. 3d). In neither condition were beads detected in EEA1 single-positive early endosomes, ruling out delayed phagocytosis.

**Fig. 3:**
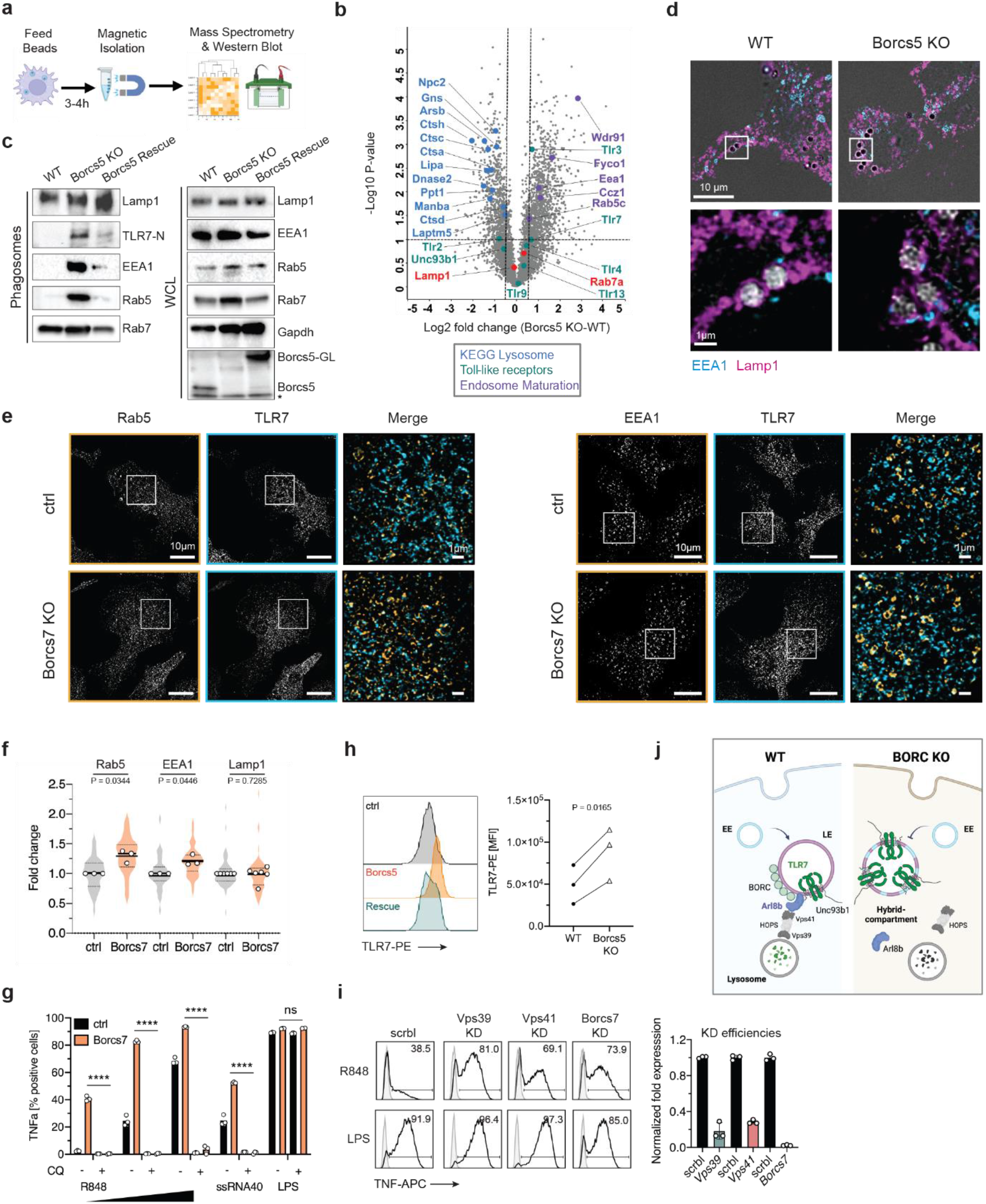
BORC controls endosome maturation and TLR7 turnover. **a**, Phagosome purification workflow. **b**, Volcano plot showing phagosome proteomics data (n=3). The horizontal and vertical lines indicate a *P*-value of 0.1 and fold change of 1.5 respectively. Selected proteins are highlighted. **c**, Western blot validation of phagosome preparations (* unspecific band). Representative data of N=3. WCL: whole cell lysate. **d**, EEA1 and Lamp1 staining of bead-containing phagosomes after 3h of feeding to Raw267.5 macrophages. Representative images of N=2. **e**, Representative images of indicated Hoxb8 macrophage lines stained for endogenous TLR7 and either Rab5 or EEA1 and imaged on a structured illumination microscope. **f**, Quantification of the colocalization between TLR7 and Rab5, EEA1, or Lamp1 in Borcs7 KO and ctrl Hoxb8 macrophages. Data is expressed as fold-change and pooled from N=3. Dots represent means of independent experiments, shaded violin shows the variation of individual cells across all experiments; ratio paired t test, two-tailed. **g**, Intracellular TNF staining of indicated Hoxb8 macrophage lines with R848 (2, 5, 10 ng/ml), ssRNA40/LyoVec (5µg/ml), or LPS (25ng/ml) in the presence or absence of chloroquine (CQ). Representative data (n=3) of N=2; unpaired t test, two-tailed. **h**, TLR7 flow staining of indicated Raw264.7 macrophage lines. Histogram shows representative raw data; dot blot shows pooled data of N=3; ratio paired t test, two-tailed. **i**, Intracellular TNF staining of indicated knock-down Hoxb8 macrophages after 8h stimulation with R848 (5ng/ml) and LPS (10ng/ml). Representative data of N=3. Right: qPCR validation of knock-down efficiencies of the presented experiment on the left. **j**, Proposed model of how BORC-Arl8b controls TLR7 receptor levels and signaling activity. EE: early endosome, LE: late endosome. \*\*\*\**P* < 0.0001.

The existence of less mature hybrid compartments would predict increased TLR7 colocalization with early endosomal markers. We performed super-resolution microscopy quantifying the degree of TLR7 colocalization with the two early endosomal markers EEA1 and Rab5 (Fig. 3e and Fig. S3c), as well as the late endosomal marker Lamp1. As predicted, there was higher overlap between TLR7 and Rab5/EEA1 in Borcs7 KO Hoxb8 macrophages, whereas colocalization with Lamp1 was unaffected (Fig. 3f). As the TLR7 distribution shifts towards less mature compartments with lower levels of major lysosomal proteases, we determined whether signaling was still dependent on acidification. Chloroquine treatment completely abrogated the TLR7 response in both WT and BORC KO macrophages (Fig. 3g). Another reported function of the BORC-Arl8b complex is the regulation of vesicle fusion between late endosomes and lysosomes^21,22^. We reasoned that by regulating lysosome fusion BORC would be able to control the turnover of membrane proteins targeted for lysosomal degradation, including TLR7. Indeed, we observed an overall accumulation of TLR7 protein in Borcs5-deficient macrophages by flow (Fig. 3h). The specificity of the anti-TLR7 antibody was validated against KO controls (Fig. S3d). To mediate the fusion between late endosomes and lysosomes, BORC-Arl8b recruits the mammalian homotypic fusion and protein sorting (HOPS) complex, a tethering complex regulating membrane fusion events with lysosomal membranes^23^. Arl8b has been shown to directly interact with the HOPS subunit VPS41^24^, and phagosomes from BORCs5 KO macrophages showed a significant depletion of VPS41 (Supplementary table 3). We therefore hypothesized that removal of HOPS complex components would recapitulate the TLR7 gain-of-function phenotype of BORC KO cells. Indeed, knock-down of the HOPS subunits Vps39 or Vps41 in Raw264.7 macrophages resulted in enhanced TLR7 signaling, providing independent confirmation on the role of impaired lysosome fusion (Fig. 3i). None of the knock-down conditions affected TLR4 signaling.

We propose a model in which the BORC-Arl8b complex safeguards the proper homeostatic turnover of TLR7. In the absence of Arl8b or BORC subunits, TLR7-containing late endosomes fuse less efficiently with lysosomes, resulting in a build-up of TLR7 within stalled hybrid endosomes (Fig. 3j).

### BORC-Arl8b interacts with TLR7 through Unc93b1

Next, we determined whether BORC-Arl8b would directly interact with nucleic acid-sensing TLRs at endosomal membranes. We reasoned that an interaction would most likely occur through Unc93b1, as it is structurally wrapped around the TLR and provides abundant binding surfaces through its long cytosolic N- and C-terminal tails^25^. We immunoprecipitated Flag-tagged Unc93b1 from Raw264.7 macrophages and observed an interaction with endogenous Arl8b (Fig. 4a). The interaction was independent of TLR7 stimulation but required functional Unc93b1 that reaches late endosomes, as the trafficking-deficient H412R mutant Unc93b1^26^ did not bind Arl8b (Fig. 4a). Alphafold multimer modeling predicted a putative interaction of Arl8b with the acidic patch (45-49 EEEEE) in the N-terminus of Unc93b1, as well as additional regions including I317 in loop 6 (Fig. 4b-c and Fig. S4b). We introduced triple alanine mutations into the predicted Unc93b1 binding surfaces including surrounding areas and showed that these mutant alleles, when expressed in Raw264.7 macrophages, phenocopied the TLR7 hyperresponse observed in Arl8b or BORC KO macrophages (Fig. S4a).

**Fig. 4:**
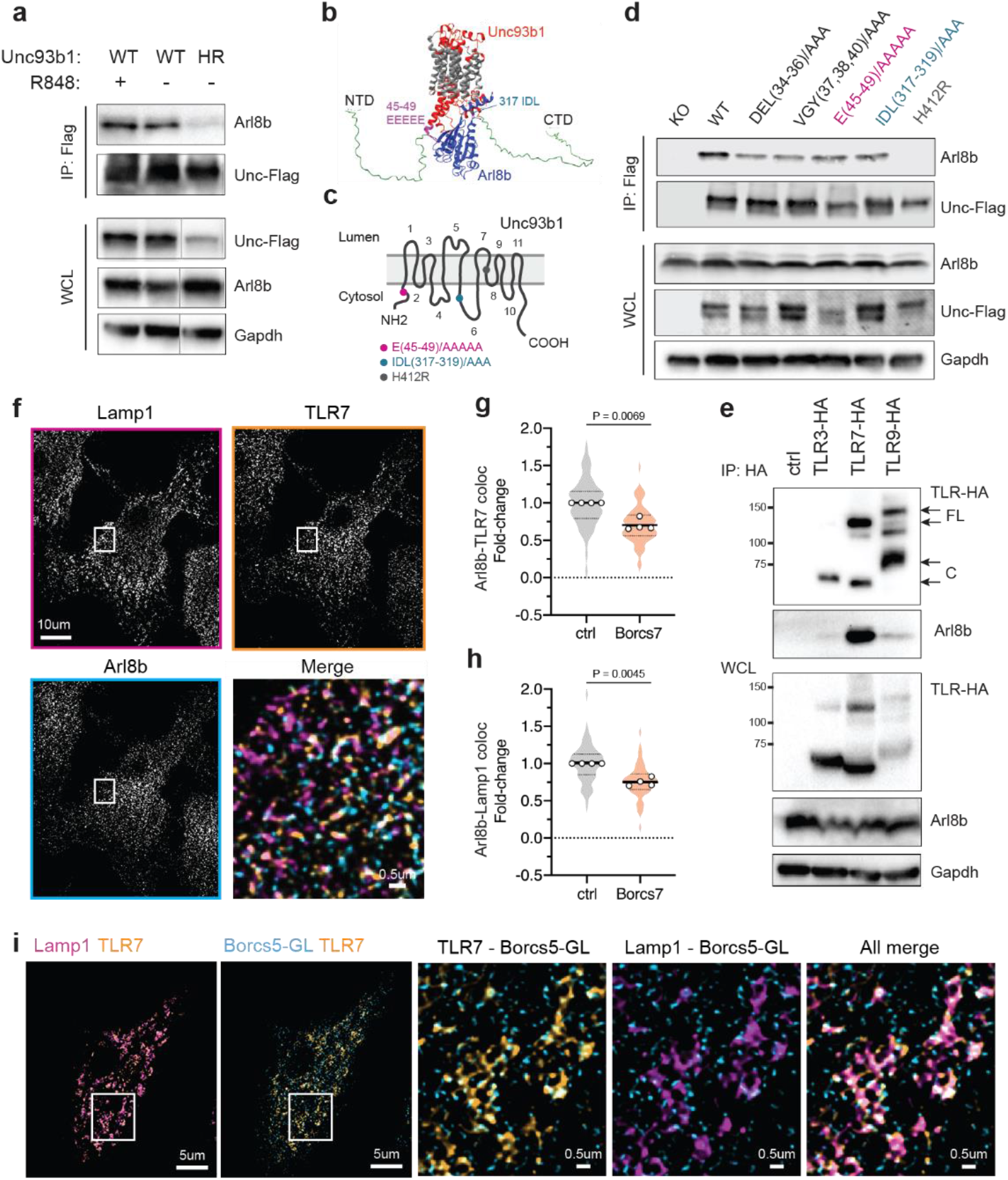
BORC-Arl8b interacts exclusively with the TLR7-Unc93b1 complex at late endosomes. **a**, Flag immunoprecipitation of Unc93b1 from Raw264.7 macrophages expressing either WT or non-functional Unc93b1^H412R^ and stimulated for 1h with R848 (200ng/ml) or left unstimulated, followed by immunoblot for Arl8b. Representative data of N=3. **b**, AlphaFold multimer modeling of Unc93b1 and Arl8b. **c**, Domain structure of Unc93b1 with highlighted mutations. **d**, Flag immunoprecipitation of Unc93b1 from Raw264.7 macrophages expressing the indicated Unc93b1 mutants and immunoblotting for Arl8b. Representative data of N=2. **e**, HA immunoprecipitation of indicated TLRs from Raw264.7 macrophages and immunoblotting for Arl8b. Representative data of N=2. **f**, WT Hoxb8 macrophages were fixed and stained with anti-Lamp1, anti-TLR7, and anti-Arl8b and imaged on a structured illumination microscope. **g**, Quantification of the colocalization between Arl8b and TLR7 in Borcs7 KO and ctrl Hoxb8 macrophages. Data is expressed as fold-change and pooled from N=4. Each dot presents the average from a single experiment, shaded violin shows the variation of individual cells across all experiments; ratio paired t test, two-tailed. **h**, Quantification of the colocalization between Arl8b and Lamp1 in Borcs7 KO and ctrl Hoxb8 macrophages. Data repeats and analysis as in **g**. **i**, Raw264.7 macrophages expressing Borcs5-GreenLantern were fixed and stained with anti-Lamp1, anti-TLR7, and anti-GFP and imaged on a structure illumination microscope.

To test whether any of these regions mediate the association with Arl8b, we performed co-immunoprecipitation experiments. All tested Unc93b1 mutations showed a clear reduction in Arl8b binding (Fig. 4d), correlating well with the TLR7 gain-of-function. One of the tested Unc93b1 alleles included the previously reported D34A mutation, which drives lupus-like disease in mice^2^; indicating a potential involvement of BORC-mediated regulation for this mutation also. As none of the individual Unc93b1 mutants showed a complete loss of Arl8b binding, contact between multiple surfaces might be required for efficient binding. After having mapped the interaction of Unc93b1 and Arl8b, we sought to determine which specific TLR-Unc93b1 complex would engage with the Arl8-BORC complex. HA-tagged TLR3, TLR7, and TLR9 were immunoprecipitated from separate Raw264.7 macrophage lines and probed for binding to Arl8b. Strikingly, only TLR7 pulled down Arl8b, whereas neither TLR3 nor 9 showed an interaction (Fig. 4e). The specific interaction with TLR7 can be explained by the fact that both TLR3 and 9 release from Unc93b1 once inside the endosome, whereas TLR7 remains associated^27^. However, how BORC still influences the signaling of TLR3 and 9 without directly interacting with the receptors will require further investigation.

To confirm the TLR7-Arl8b interaction at late endosomal membranes, we performed super-resolution imaging of cells labeled for endogenous TLR7, Arl8b, and Lamp1 (Fig. 4f). Antibody specificities against mouse TLR7 and Arl8b were confirmed using the respective KO lines (Fig. S4c). In the absence of BORC, colocalization between Arl8b and TLR7 decreased (Fig. 4g), due to reduced recruitment of Arl8b to late endosomes (Fig. 4h and Fig. S4d). Using Raw264.7 macrophages expressing Borcs5-GreenLantern, we also showed that TLR7 colocalizes with BORC at late endosomes (Fig. 4i), although in Co-IP experiments we had not been able to directly pull down BORC subunits with TLR7.

Taken together, our findings point to dual roles for BORC, which are 1) mediating the fusion of late endosomes and lysosomes and 2) recognizing TLR7-Unc93b1 as specific cargo to be sorted into these lysosomes. To uncouple these two functions, we assessed endosome maturation in macrophage lines with a compromised Unc93b1-Arl8b interaction, but otherwise harboring an intact BORC complex. As expected, endosome maturation was normal in cells expressing Unc93b1 mutations with reduced Arl8b interaction, indicated by the absence of early endosome markers on phagosome preparations (Fig. S4e). Our data supports a model in which the Arl8b-Unc93b1 interaction is crucial to recognize TLR7 as specific cargo for lysosomal sorting to maintain healthy TLR7 levels.

### Breaking the interaction between Arl8b and Unc93b1 causes human lupus

To confirm the relevance of our findings for human physiology, we analyzed patients with autoimmune disease and identified a young female with systemic lupus erythematosus (SLE) and a monoallelic UNC93B1 mutation (Fig. 5a). The variant was inherited from her father, who had not yet come to medical attention (Fig. 5a). The mutant allele p.Glu49dup carried a glutamic acid insertion within the N-terminal 5xE acidic patch flanking the ARL8B binding motif thereby extending it to 6xE (Fig. 5a-b and Fig. S5a). UNC93B1 6xE has not been reported in public databases (GnomAD, ExAC, and GME) or in our in-house exome database (5,600 exomes) and thus is a private variant. Phylogenetic analysis of 420 homologous UNC93B1 vertebrate protein sequences encompassing 116 mammalian sequences showed high conservation of the 5xE stretch (Fig. 5c and Fig. S5b). No other structural or genetic gene variants associated with SLE^28^ were identified. UNC93B1 protein levels were comparable in PBMCs from the patient and her parents (Fig. 5d) indicating that the variant did not alter protein expression.

**Fig. 5:**
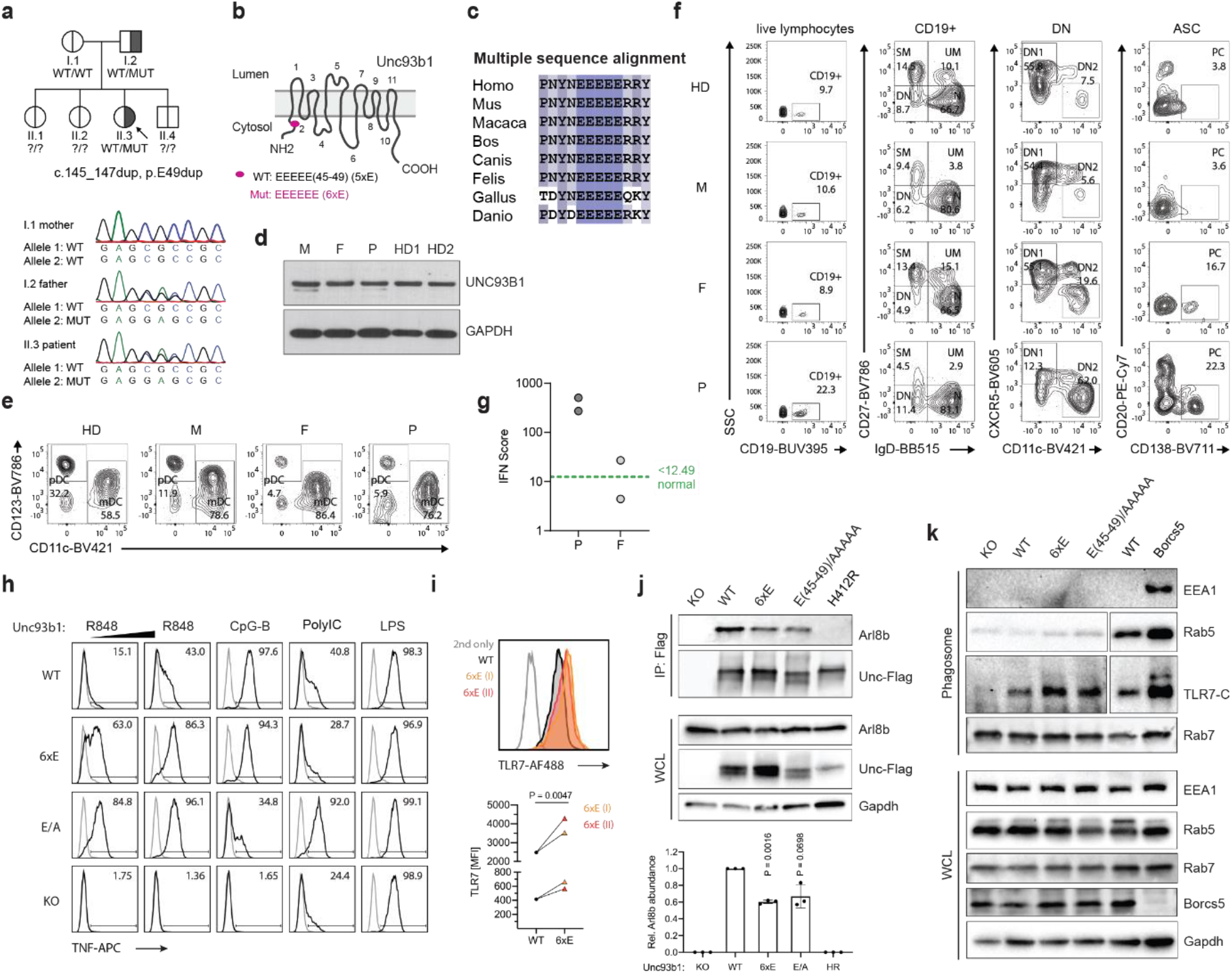
Lupus patient with an Unc93b1 mutation in the Arl8b-binding domain. **a,** Family pedigree with affected child and Sanger sequencing electropherograms from PBMCs genomic DNA of the patient. **b,** Domain structure of Unc93b1 with highlighted patient mutation. **c,** Multiple sequence alignment of the Unc93b1 5xE region. **d,** UNC93B1 immunoblot from PBMC total protein extracts. **e-f,** Flow cytometric immune phenotype of peripheral blood from a healthy donor (HD), the patient’s mother (M), father (F), and the patient (P). SM: switched memory, UM: unswitched memory, N: naive, DN: double negative, ASC: antibody secreting cell, PC: plasma cell, mDC: myeloid dendritic cells, pDC: plasmacytoid dendritic cells. **g,** IFN signatures in PBMCs of patient (P) and father (F). An IFN score of 12.49 (dashed green line) indicates the median IFN score of 10 healthy controls plus 2 SD. **h,** Intracellular TNF staining of Raw264.7 macrophages expressing the indicated Unc93b1 variants with R848 (5 and 10 ng/ml), CpG-B (300 nM), PolyIC (10µg/ml) or LPS (25ng/ml). Representative data of N=2. **i**, Representative TLR7 flow staining of indicated Raw264.7 macrophage lines. Dot plot shows pooled data from N=2; ratio paired t test, two-tailed. **j**, Flag immunoprecipitation of Unc93b1 from Raw264.7 macrophages expressing the indicated Unc93b1 mutants and immunoblotting for Arl8b. Representative data of N=3. WCL: whole cell lysate. Bar graph shows the quantification of all N=3; ratio paired t test, two-tailed. **k**, Western blot of phagosome preparations. Representative data of N=2. WCL: whole cell lysate.

The patient developed progressive autoimmune symptoms at the age of 7, including atopic dermatitis, Hashimotos’s thyroiditis, sialadenits, generalized lymphadenopathy, and splenomegaly (Supplementary Data 1). At the time of evaluation, the patient exhibited antinuclear antibodies (ANA), leucopenia, massive hypergammaglobulinemia, and complement consumption (Supplementary Data 2 and Supplementary Table 1). Deep immune phenotyping of PBMCs showed accumulation of total CD19^+^ B-cells, IgD^-^CD27^-^ double-negative (DN) B-cells and their CXCR5^-^CD11c^+^ DN2 subset [i.e., age-associated B-cells (ABC)], and CD20^-^CD138^+^ plasma cells in the patient and to some extend in her father being compatible with increased TLR7 signaling (Fig. 5e)^29^. CD24^+^CD38^+^ transitional B-cells and their CD10^+^CD38^+^ transitional 1 subset, total CD19^+^ B- and naïve CD21^low^CD38^low^ activated B-cells were increased in the patient (Fig. S5c). In addition, CD4^+^/CD8^+^ T-cell ratio, CD4^+^ and CD8^+^CD38^+^HLA-DR^+^ activated T-cells, and CD8^+^CD57^+^PD-1^+^ senescent T-cells were increased in the patient and to some extend in her father reflecting reactive T-cell states. In the patient, CD4^+^CD45R0^+^CCCR6^-^ helper T-cells and their CCR4^+^CXCR3^-^ helper type 2 (Th2) was decreased reflecting lupus disease activity (Fig. S5d)^30,31^. CD56^+^CD57^+^ mature NK-cells were reduced and CD16^+^HLA-DR^+^ pro-inflammatory monocytes were increased in the patient (Fig. 5f), while lin^-^CD123^+^ pDCs were reduced in the patient and her father reflecting an inflammatory state (Fig. 5f)^31,32^. In line with that, the patient had a highly increased interferon signature, while her father showed intermediate levels (Fig. 5g)^33^. These findings prompted us to further investigate the patient 6xE allele in Raw264.7 macrophages. In response to R848 stimulation, TNF production as well as *Ifnb* induction was significantly stronger in cells carrying the 6xE variant compared to cells expressing the wild type allele (Fig. 5h and Fig. S5f). No differences were observed for TLR9 or TLR3 signaling (Fig. 5h). Unc93b1 KO cells served as negative control. Notably, cells expressing the E/A Unc93b1 mutation showed an even greater response to TLR stimulation, suggesting that the alanine-replacement has a broader functional effect on Unc93b1 than the single E insertion (Fig. 5h). Equal expression levels of the various Unc93b1 mutant variants were validated by flow (Fig. S5g). Consistent with our findings in BORC KO cells, we saw an accumulation of TLR7 protein in two independently-derived macrophage lines expressing the patient allele (Fig. 5i), without changes at the *Tlr7* transcript level (Fig. S5h). As expected, the additional E insertion reduced the binding between Unc93b1 and Arl8b (Fig. 5j), indicating that the patient allele diminishes BORC-mediated TLR7 degradative sorting into lysosomes, leading to receptor accumulation and increased signaling. We confirmed an accumulation of cleaved TLR7 in phagosomes of 6xE Unc93b1-expressing macrophages (Fig. 5k). As the BORC complex itself was not compromised in these cells, endosome maturation proceeded otherwise normal (Fig. 5k). Our findings demonstrate that interfering with the BORC-mediated homeostatic TLR7 turnover can lead to SLE in humans.

## Discussion

Our work establishes the importance of an intact endomembrane system supporting a healthy homeostatic turnover of TLR7 to prevent autoimmune disease. As TLR7 expression levels must be tightly controlled to avoid excessive stimulation^1,4,5,34^, mechanisms stabilizing TLR7 protein would be likewise expected to break self-tolerance.

A Glu49 duplication in Unc93b1, identified in a young lupus patient, weakens the interaction with the BORC-Arl8b complex resulting in reduced degradative sorting and accumulation of TLR7 in late endosomes. Even though the monoallelic variant clearly drives TLR7 gain-of-function, disease penetrance is incomplete, demonstrated by the absence of symptoms in the father, who is also a carrier. Incomplete disease penetrance is common for lupus-associated risk alleles, as also environmental and (epi)genetic factors are known to contribute^35^. Recently, additional families with UNC93B1 mutations in distinct protein regions have been described that also drive lupus disease^7^. Taken together, these two reports establish a key role for Unc93b1 in preventing excessive TLR7 signaling and autoimmunity in humans. The involvement of the BORC complex in immune regulation is a new finding, and might have been obscured by the central role BORC plays in maintaining neuronal integrity. Despite being expressed ubiquitously, the BORC-Arl8b complex is particularly important in long-range vesicle transport within axons^36^. As a result, mice with attenuated BORC function develop progressive axonal dystrophy and impaired motor function^37^. Full-body KO mice of individual BORC subunits or Arl8b are not viable^21,37–39^. These animal studies give reason to predict that humans with attenuated BORC function would foremost present with neurodegenerative disorders; which notably, quite frequently cooccur with autoinflammatory or autoimmune reactions and often share genetic variants and pathways^40^. This might also explain why no BORC mutations have so far been associated with autoimmune diseases in GWAS studies.

The BORC-Arl8b complex facilitates multiple cellular functions and we show here that not all cell types utilize these functions equally. In Hela cells, BORC regulates lysosome positioning and autophagy^12,19^, whereas in macrophages its main role appears to be sorting of endocytic and biosynthetic cargo into lysosomes. The same function seems to apply to dendritic cells, in which Arl8b-deficiency delays lysosomal maturation and cargo delivery to lysosomes, resulting in reduced microbial killing and antigen presentation^22^. In TLR7-stimulated plasmacytoid dendritic cells, Arl8b controls the trafficking of TLR7-containing vesicles into the cell periphery for type I interferon induction^41^, a phenomenon we do not see in macrophages.

Our work provides the first example of how endosomal dysregulation can result in TLR7 gain-of-function and lupus disease in humans. It expands the repertoire of cellular mechanisms important to restrict pathological TLR7 activity and paves the way for improved treatment through TLR7-targeted precision medicine.

## Methods

### Patients

All subjects and/or their legal guardians gave written informed consent for immunological and genetic investigations that were carried out with approval from the institutional review board of the Ludwig-Maximilians-Universität München (project number 66-14).

### Antibodies and reagents

The following antibodies were used for immunoblots and immunoprecipitations: anti-HA as purified antibody or matrix (3F10, Roche), anti-FLAG as purified antibody or matrix (M2, Sigma), anti-Unc93b1 (1:1000, PA5–20510, Thermo Scientific), anti-GAPDH (1:1000, MA5-15738, Invitrogen), anti-mLamp-1 (1:1000, AF4320, R&D Systems), anti-Phospho-p38 (1:1000, D3F9, 4511T, Cell Signaling), anti-p38 (1:1000, D13E1, 8690T, Cell Signaling), anti-Phospho-p44/42 (ERK1/2) (1:1000, 197G2, 4377T, Cell Signaling), anti-p44/42 (ERK1/2) (1:1000, 137F5, Cell Signaling), anti-IκBα (1:1000, L35A5, 4814, Cell Signaling), anti-Borcs5 (1:500, 17169-1-AP, Proteintech), anti-Borcs7 (1:500, PAB23142, Abnova), anti-Arl8b (1:500, 13049-1-AP, Proteintech), anti-TLR7 (1:500, MAB7156, R&D systems), anti-EEA1 (1:1000, clone C45B10, 3288S, Cell Signaling), anti-LC3b (1:1000,clone D11,3868,Cell Signaling), anti-Rab7 (1:1000, D95F2, 9367, Cell Signaling), anti-Rab5 (1:1000, C8B1, 3547, Cell Signaling). Secondary antibodies: AlexaFluor 680 anti-mouse IgG (A21057, Invitrogen), AlexaFluor 680 anti-rabbit IgG (A21076, Invitrogen), anti-Rabbit HRP (111-035-144, Jackson ImmunoResearch), anti-Mouse HRP (115-035-166, Jackson ImmunoResearch), anti-Rat HRP (112-035-062, Jackson ImmunoResearch).

Antibodies and reagents used for flow cytometry were: anti-Lamp1 FITC (1:200, 553793, BD Pharmingen) anti-TNFα (1:200, MP6-XT22, eBioscience), purified anti-CD16/32 Fc Block (1:200, 2.4G2), anti-TLR7 PE (1:200, clone A94B10, 160003, Biolegend), eFluor 780 (1:1000, 65-0865-14, Invitrogen), eFluor506 (1:500, 65-0866-14, Invitrogen). Cells were fed with DQ-BSA Red (10µg/ml, D12051, Invitrogen), Dextran-AF680 (50µg/ml, MW3000, D34681, Invitrogen), Dextran-AF647 (50µg/ml, D22914, Invitrogen) Bafilomycin A (100nM, B1793, Sigma), Chloroquine (50µM, tlrl-chq, Invivogen).

Anti-human antibodies and reagents for flow cytometry were: BV480-anti-CD45 (HI30), BUV496-anti-CD3 (UCHT-1), BUV395-anti-CD4 (RPA-T4), PE-Cy5-anti-CD8 (RPA-T8), BUV737anti-CD45RA (HI100), BB515-anti-CD45R0 (UCHL1), APC-R700-anti-CD27 (M-T271), BV711-anti-HLA-DR (G46-6), PE-CF594-anti-CXCR3 (1C6), BUV395-anti-CD19 (SJ25C1), PE-Cy7-anti-CD20 (2H7), BB515-anti-IgD (IA6-2), BUV496-anti-IgM (UCH-B1), BUV737-anti-CD21 (B-Ly4), BV711-anti-CD138 (MI15), BB700-anti-CD24 (ML5), PE-anti-CD10 (HI10a), BUV737-anti-CD8 (SK1), R718-anti-CD19 (SJ25C1), BV605-anti-PD-1 (EH12.1), PE-CF594-anti-CD56 (NCAM16.2), PE-BB515-anti-CD57 (NK-1), BB700-anti-CD14 (M5E2) and Fixable Viability Stain 780 all from BD and APC-anti-CD127 (A019D5), BV421-anti-CCR7 (G043H7), BV785-anti-CCR6 (G034E3), PE-Cy7-anti-CCR4 (L291H4), BV605-anti-CCR4 (J252D4), BV650-anti-CD38 (HB-7), BV786-anti-CD27 (323), BV785-anti-CD123 (6H6), BV421-anti-CD11c (Bu15), APC-anti-CD16 (3G8), PE-Cy7-anti-CD33 (P67.6) from BioLegend, San Diego, USA.

Antibodies and dyes used for Immunofluorescence were : 1) Primary antibodies :anti-Arl8b (1:200, 13049-1-AP, Proteintech), anti-TLR7 (1:200, clone A94B10, MABF2273,Sigma), anti-EEA1 (1:200, clone C45B10, 3288S, Cell Signaling), anti-GFP (1:200, 11814460001, Roche), anti-GFP (1:500, G10362, Invitrogen), anti-Rab7 (1:100, D95F2, 9367, Cell Signaling), anti-Rab5(1:200, C8B1, 3547, Cell Signaling), anti-Rab11 (1:50, D4F5, 5589, Cell Signaling), anti-Lamp1(1:800, AF4320, R&D),anti-Lamp1(1:200, AB24170, Abcam),anti-Lamp1(1:200, 1D4B, 553792, BD Pharmingen). 2) Secondary antibodies: anti-mouse-488 (1:1000, A11029, Invitrogen), anti-rabbit-488 (1:1000, A21206, Invitrogen), anti-rabbit-568 (1:1000, A10042, Invitrogen), anti-goat-647 (1:1000, A32849, Invitrogen). 3) Dyes: NucBlue Live Cell Stain (2 drops/ml, R37605, Molecular Probes), phalloidin-568 (A12380, Invitrogen), DAPI.

### TLR ligands

CpG-B (ODN1668: TCCATGACGTTCCTGATGCT, all phosphorothioate linkages) and ssRNA40 (5’-GCCCGUCUGUUGUGUGACUC-3’, all phosphothioate linkages) were synthesized by Integrated DNA Technologies. ssRNA40 was complexed with LyoVec (lyec-1, Invivogen) for 30min at RT before stimulation according to manufacturer’s instructions. R848, PolyIC (HMW), Pam3CSK4, and LPS-EB (*E. coli* 0111:B4) were purchased from InvivoGen.

### Cell and tissue culture conditions

HeLa (DSMZ collection, Germany), HEK293T (ATCC collection) and GP2–293 packaging cell lines (Clontech) were cultured in DMEM (10938-025, Invitrogen) complete media supplemented with 10% (vol/vol) FCS (F0804, Sigma), 1% penicillin-streptomycin (15140122, Gibco), 1% sodium pyruvate (11360070, Gibco), 1% L-glutamine (25030081, Gibco) and 1% HEPES (15630056, Gibco) at 37°C and 5% CO_2_. Unc93b1 KO Raw264.7 macrophages (described in ^42^) and Cas9-expressing Hoxb8 progenitors (described in ^43^) were a kind gift from the Barton Lab, UC Berkeley. THP-1 cells and WT Raw264.7 cells were a kind gift from the Zychlinsky Lab, MPI for Infection Biology. RAW264.7 and THP-1 cell lines were cultured in RPMI 1640 (31860-025, Gibco) using the same supplements as listed above. The myeloid Hoxb8 progenitors were cultivated in RPMI 1640 medium with 10 % FCS, 1 % L-Glutamine, 1 % Sodium-Pyruvate, 1 % HEPES, 1 % Penicillin/Streptomycin, 2 % GM-CSF-conditioned medium produced by a B16 murine melanoma cell line, 0,0003 % beta-mercaptoethanol (21985-023, Gibco) and 0,02 % beta-estradiol (Sigma).

PBMCs were isolated by Ficoll-Hypaque density gradient centrifugation (Biochrom, Berlin, Germany) and cultured in RPMI1640 supplemented with 2 mM Glutamax, 100 U/ml penicillin, 100 µg/ml streptomycin, and 10 % FCS (Thermo Fisher Scientific, Waltham, USA) at 37°C and 5 % CO_2_.

### Generation and differentiation of Hoxb8 macrophages

For a detailed description of how to generate Hoxb8 progenitors and differentiate them into macrophages see ^43,44^. In brief, for the generation of Hoxb8 progenitors, mouse bone marrow was flushed and cultured for 2 days in stem cell medium (DMEM + 15% FCS + 25ng/ml SCF (PeproTech) + 10ng/ml IL-3 (PeproTech) + 20ng/ml IL-6 (PeproTech)). Cells were spinfected with retrovirus containing the FLAG-ER-Hoxb8-MSCV-Neo construct and selected for survival/immortalization during subsequent passages in progenitor medium containing beta-estradiol. For the differentiation of Hoxb8 progenitors into macrophages, the progenitor medium was removed by centrifugation at 1200 rpm for 5min, then the cells were washed twice with 1xPBS. 1 x10^6^ progenitor cells were plated in 10ml macrophage media (RPMI 1640 medium, 10 % fetal calf serum, 1 % L-Glutamine, 1 % Sodium-Pyruvate, 1 % HEPES, 1 % Penicillin/Streptomycin, 10 % conditioned M-CSF medium produced by a 3T3 mouse embryonic fibroblast cell line, and 0,0003 % β-mercapto-ethanol) in a non-TC treated 10cm petri-dish. On day 3, 10ml fresh macrophage media was added. On day 6, 10ml of the media was exchanged with fresh 10 ml macrophage media. On day 7-9, the macrophages were ready to be used for experiments.

### THP-1 differentiation PMA

THP-1 cells were differentiated for 48 hours using 25 nM PMA (tlrl-pma, Invivogen) in RPMI media supplemented as mentioned above. After 48 hours, the media was removed and cells were washed once with RPMI supplemented media and replaced with fresh media for another 24 hours before stimulation experiments.

### Retro and lentiviral transduction

For viral transduction, VSV-G-pseudotyped retrovirus was produced in GP2–293 packaging cells (Clontech) and VSV-G-pseudotyped lentivirus, in HEK–293T cells. GP2–293 or HEK-293T cells were transfected with either retroviral vectors and pVSV-G or lentiviral vectors, pVSV-G and pPax2 using Lipofectamine 3000 (L300001, Invitrogen) reagent. The day after transfection, packaging cells were moved to 32°C for 24h. 2 days post transfection, viral supernatant was collected and cleared by spinning twice at 2000rpm for 5min. The cleared supernatant (with polybrene at final 5μg/ml) was used to infect recipient cells. For transduction, cells were spin-infected at 1000g for 30min. Infected cells were incubated overnight at 32°C and protein expression was checked 48h later. Recipient cells were either sorted on fluorescent reporters (BD FACS Aria II Floyd Sorter) or drug-selected with Puromycin (A11138-03, Gibco) starting 48h after transduction.

### CRSPR/Cas9 KO in Hoxb8 progenitors

For a detailed description see ^43^. In brief, Cas9-expressing Hoxb8 progenitors were retrovirally transduced with pLentiGuide Puro (Addgene #52963) in which the gRNA sequences have been cloned using the BsmBI restriction site. gRNA sequences were generated through annealing of two complementary primers with the respective overhangs. 48h after spinfection cells were selected with Puromycin in progenitor medium. Used gRNA sequences are listed in the key resource table.

### CRISPR RNP Knock-out

Crisprmax Transfection Reagent (CMAX00008, Invitrogen) was used to generate RAW264.7 BORC subunit knock-out cell lines. 1.5µmol of guide RNA oligos and 1.5µmol of Alt-R® S.p. Cas9 Nuclease v3 protein (1081058, IDT) were mixed with 0.6µl of Cas9 PLUS reagent in Optimem media to assemble the RNP complexes in a volume of 25µl. The assembled complexes were mixed with 1.2µl of CRISPRMAX transfection reagent diluted in 25µl Optimem and incubated for 20mins at RT. RNP complexes (10nM final) were transfected into 40,000 cells in a 96-well plastic U-bottom plate. 72 hours post-transfection, cells were single-cell sorted for screening individual knockout clones by western blot.

### Immunofluorescence

Cells were plated onto glass coverslips (No.15H) and allowed to settle overnight. Cells were washed 1x with PBS, fixed with 2% PFA-PBS for 15min at RT, and permeabilized with 0.5% saponin (Roth)-PBS for 5min at RT. To quench PFA autofluorescence coverslips were treated with ammonium chloride 50mM/0.1% saponin-PBS for 10min at RT. After washing 3x with PBS, cells were blocked in 1% BSA/0.1% saponin-PBS with 4% horse serum (26050-070, Gibco) for 1h at RT. Slides were stained in blocking buffer with primary antibodies overnight at 4°C, washed 3x with PBS and incubated for 1h with secondary antibodies in blocking buffer at RT. Cells were washed 3x in PBS and mounted using ProLong Glass antifade mountant (P36982, Invitrogen). Cells were imaged on a Zeiss Elyra 7 with lattice SIM with a 63X oil immersion objective.

### Superresolution SIM microscopy

Structured Illumination microscopy was performed on the Zeiss Elyra 7 lattice SIM microscope equipped with 405, 488, 561 and 642 nm laser for excitation. Z-stacks from fixed cells were acquired on the SIM mode using a 63X/1.6 oil immersion objective and pco.edge 4.2 sCMOS camera. Raw images were SIM processed and channel aligned using Zeiss default settings in Zen Black. A new channel alignment was acquired for every imaging session, using Hoxb8 macrophages triple-stained for Lamp1 (in AlexaFluor 488, 568, and 647). The calculated off-set from this image was applied to all other images for channel alignment to correct for the shift in xyz between different channels. The completed super-resolution Z-series was visualized and analyzed using Fiji (ImageJ). For image quantification in Cell Profiler individual slices were used. Images for testing antibody specificity were acquired using the wide-field mode. For each new staining panel bleed-through controls were performed with single-stained cells.

### Ratiometric fluorescence microscopy

Ratiometric endosomal pH measurements were performed as previously described in ^45^. In brief, VWR confocal dishes (734-2906) were coated with 10µg fibronectin (F1141, Sigma) on the Ø20 mm glass bottom center, overnight at 37°C. Cells were seeded on freshly-coated plates and loaded with 0.37mg/mL of FITC-Dextran (Fixable, MW10000, D1820, Invitrogen) and 0.075mg/ml Alexa 647-Dextran (MW10000, D22914, Invitrogen) overnight. The next day, cells were washed in 1xPBS and incubated with CellMask Orange (1:2000, C10045, Invitrogen) for 10mins at 37°C. Cells were washed again in PBS, taken up in fluoroBrite imaging media (A1896701, Gibco) with 10% FCS + 1 % L-Glutamine and imaged on a spinning disk confocal. Confocal microscopy was performed on a Nikon Eclipse Ti2 inverted microscope attached to an Andor Dragonfly 200 spinning disk unit with Andor Zyla sCMOS camera. The microscope was equipped with 405, 445, 488, 514, 561 and 637nm lasers and operated by Fusion software 2.3. Samples were imaged live using an Apo TIRF 60X/1.49 Oil objective (MRD01691).

### Endosome positioning

HeLa cells were plated on 6-well plates and transfected the next day with either 1) Arl8b expression plasmid or empty vector control using Lipofectamine 3000 or 2) siRNA against Borcs5 or scrambled siRNA according to protocol ‘siRNA knock down’. After 48h cells were fixed and stained with anti-Lamp1, phalloidin-568 (A12380, Invitrogen), and DAPI according to protocol ‘Immunofluorescence’. Phalloidin and DAPI served to visualize cell border and nuclei for automated quantification.

Hoxb8 macrophage lines were fixed and stained the same way. Cells were imaged on a standard spinning disk confocal or laser scanning confocal.

For live-cell imaging, RAW264.7 cells were seeded on fibronectin coated 8-well chamber slides overnight. Next day, the cells were loaded with CellMask^TM^Orange plasma membrane stain (1:2000), LysoTracker^TM^ Deep Red (1:40000, L12492, Invitrogen) and NucBlue Live Cell Stain diluted in FluoroBrite imaging media (A1896701, Gibco) with 10% FCS + 1% Glutamine for live cell imaging.

### Immunoprecipitation and western blot analysis

For Flag-IP: Cells from one confluent 15cm plate were washed in ice-cooled PBS, scraped and lysed in 1% digitonin lysis buffer (20 mM Tris/HCl pH 7.4, 150 mM NaCl, 1 mM CaCl2, 1 mM MgCl2, 10% Glycerol, 1 × Complete EDTA-free cocktail (11836170001, Roche), PhosSTOP phosphatase inhibitor (04906845001, Roche), 1mM PMSF (10837091001, Roche) and GTP (2mM, G8877, Sigma). After a 1h incubation at 4°C on a rotator, lysates were cleared of insoluble material by centrifugation. For immunoprecipitation, lysates were incubated with anti-FLAG matrix (M8823, Millipore; pre-blocked with 1% BSA-PBS for 1h) for 2h, washed four times in 0.1% digitonin wash buffer (20 mM Tris/HCl pH 7.4, 150 mM NaCl, 1 mM CaCl2, 1 mM MgCl2, 10% Glycerol), and precipitated proteins competitively eluted with Flag peptide (150 ng/µl; F4799, Sigma). Lysates were denatured in SDS-PAGE buffer (4x Lämmli buffer (Bio-Rad) + 20mM DTT) at room temperature for 1h. Proteins were separated by SDS-PAGE (Bio-Rad TGX precast gels) and transferred to Immobilon PVDF membranes (Millipore) in a Trans-Blot Turbo transfer system (Bio-Rad). Membranes were probed with the indicated antibodies and developed using the Licor scanning System (for quantitative western) or the Pierce ECL Plus Western Blotting Substrate (32132, Thermo) and Bio-Rad ChemiDoc imaging system. Samples were normalized based on equal bait protein in eluates, or Gapdh in whole cell lysates.

HA-IPs were performed in the same way, except cleared lysates were incubated with anti-HA matrix (11815016001, Roche) (pre-blocked with 1% BSA-PBS for 1h) for 2h at 4°C and washed four times and precipitated protein were competitively eluted with HA-peptide (100ng/ml; I2149, Sigma).

### Flow Cytometry

Mouse macrophages were seeded into non-TC treated round-bottom 96-well plates. The next day cells were stimulated with the indicated TLR ligands. To measure TNFα production, BrefeldinA (BD GolgiPlug) was added to cells 30min after stimulation, and cells were collected after an additional 5.5h. Dead cells were stained with eFluor 506 or 780 fixable live/dead dye. Cells were blocked with Fc Block anti-Mouse CD16/CD32 (1:200, 101301, Biolegend) for 10min at RT and then stained for intracellular TNFα with the BD Fixation & Permeabilization kit according to the manufacturer’s instruction. Data were acquired on a LSRFortessa or CytoFLEX (Beckman Coulter).

For immune phenotyping, whole blood was antibody stained in brilliant stain Buffer (Becton Dickinson (BD), San Jose, USA) at ambient temperature, lysed with BD Lyse-fix buffer, washed 2 times and live cells were acquired on a BD LSR Fortessa cytometer with FACS Diva software.

### Phagosome preparation

Raw264.7 cells in a confluent 15cm dish were incubated with ∼10^8^ magnetic beads (1μm size, Polysciences) for 4h. After rigorous washing in PBS, cells were scraped in 10ml sucrose homogenization buffer (SHB: 250mM sucrose, 3mM imidazole pH 7.4) and pelleted by centrifugation. Cells were resuspended in 2ml SHB plus protease inhibitor cocktail (Roche) and 1mM PMSF and disrupted by 20 strokes in a steel dounce homogenizer. The disrupted cells were gently rocked for 10min on ice to free endosomes. Beads were collected with a magnet (Dynal) and washed 4x with SHB plus PMSF. After the final wash, phagosome preparations were denatured in 2x SDS-PAGE buffer (4xLämmli buffer + 20mM DTT) at room temperature for 1h and analyzed by western blot.

For proteomic analysis, bead-containing phagosomes were additionally washed 4x with ice-cold 100mM ammonium bicarbonate and snap frozen.

### ELISA

Cells were seeded into tissue culture-treated flat-bottom 96-well plates. The next day, cells were stimulated with the indicated TLR ligands for either 8 hours (Raw264.7 Mphs) or overnight (THP-1). Supernatants were analyzed using the BioLegend ELISA MAX™ Deluxe Set Mouse TNF-α kit or Human IL-8 kit according to the manufacturer’s instructions.

### siRNA knock down

Hela cells were plated in a 6-well format. 0.5µg Plasmid DNA and 2µM siRNA were diluted in Optimem (3198-062, Gibco) and mixed. 3µl DharmaFectDuo (T-2010) was diluted in Optimem and incubated for 5min at RT. DNA/siRNA mixture was added to the diluted DharmaFectDuo and incubated 20min at RT. The cells were transfected with the complexes and used for assays 48h post-transfection. siRNAs were purchased from Dharmacon and are listed in the key resource table.

For THP-1 and Raw246.7 cells, 1.5µmol siRNA and 0.5µl Cas9 PLUS reagent were diluted in Optimem. The siRNA was mixed with 0.6µl of Crisprmax Transfection Reagent (CMAX00008, Invitrogen) diluted in Optimem and incubated for 10min at RT. 10^5^ THP-1 cells or 4x 10^4^ RAW246.7 cells in 100µl were added to the 50µl transfection mix in a flat bottom 96well (non-TC treated) plate. THP-1 cells were differentiated simultaneously as described earlier. Cells were stimulated 72h post-transfection.

### qPCR

Cells from a confluent 6-well were lysed and RNA was purified using the NucleoSpin RNA Plus kit for RNA purification (740984.50, Macherey-Nagel) according to manufacturer’s instructions. cDNA was prepared from 100–500ng RNA with iScript cDNA synthesis kit (1708891, Bio-Rad), and quantitative PCR was performed with iTaq Universal SYBR Green Supermix (1725121, Bio-Rad) on a StepOnePlus thermocycler (Applied Biosystems).

Normalization of gene expression was performed to the two reference genes *Gapdh* and *Hprt* through geometric averaging^46^. Primers were synthesized by IDT and are listed in the key resource table.

### Site-directed mutagenesis

Site-directed mutagenesis of Unc93b1 was performed with the Q5 Site Directed Mutagenesis Kit (E0552S, New England Biolabs) according to the manufacturer’s instructions.

### Protein modelling

AlphaFold multimer was used via ColabFold to predict the protein multimer model of human UNC93B1 with human ARL8B. The best prediction out of the five predicted models was chosen for further examination. The contacts were mapped using ChimeraX. The best model was compared to the Cryo-EM model of TLR7 in complex with UNC93B1^47^ using ChimeraX.

### Statistics

Statistical parameters, including the exact value of n and statistical significance, are reported in the Figures and Figure Legends, whereby n refers to the number of repeats within the same experiment. Representative images have been repeated at least three times, unless otherwise stated in the figure legends. Data is judged to be statistically significant when p < 0.05 by Student’s t-test. For experiments with two groups Two-tailed *t*-test, paired or unpaired was used. To compare means of different groups across a dose response, a two-way ANOVA followed by a Bonferroni posttest was used. In figures, asterisks denote statistical significance (*, p < 0.05; **, p < 0.01; ***, p < 0.001). Statistical analysis was performed in GraphPad PRISM 9 (Graph Pad *Software* Inc.).

Additional methods can be found in the Supplementary information.

## Acknowledgements

We thank the patient and her family for participation in the study. We acknowledge the contribution of the clinical (Veit Grote, MD; Anna-Lisa Lanz, MD; Bärbel Lange-Sperandio, MD), radiological (Marco Paolini, MD), immunodiagnostic (Irmgard Eckerlein; Mayumi Hofmann; Eva Eisl, Raffaele Conca), and genetic (Wendy Aloo; Meino Rohlfs, PhD) teams at the Dr. von Hauner Children’s Hospital and of the clinical chemistry team at the Institute of Laboratory Medicine (Peter Eichhorn, MD) both at the University Hospital, LMU Munich, Germany. We thank Noemie Quinson, Ioulia Sampani, Julian Bünting, and Daniel Landsem for assistance with experiments; Sanket Gosavi for protein modeling; the Flow Cytometry Core Facility (FCCF) of the German Rheumatism Research Center Berlin (DRFZ) for assistance with cell sorting; Stefan Bauer (University of Marburg, Germany) for sharing bone marrow of TLR7 KO mice; Beata Lukaszewska-McGreal for proteome sample preparation. Thanks to Arturo Zychlinsky and Greg Barton for carefully reading the manuscript. Figures were created with BioRender.com.

This work was funded by the Max Planck Society (D.M. and O.M.), the German Research Foundation (DFG) (679160 to O.M.), a Marie Curie fellowship from the European Research Council (841440 to O.M.), a PhD fellowship from the International Max Planck Research School (IMPRS to H.M.). F. H. was funded by the Care-for-Rare Foundation (C4R, 160073), the Else Kröner-Fresenius Stiftung (EKFS, 2017_A110), and the German Federal Ministry of Education and Research (BMBF) through a grant to the German genetic multi-organ Auto-Immunity Network (GAIN, grant code 01GM2206D). This work was further supported by grants from the Deutsche Forschungsgemeinschaft (DFG; KFO249 160548243, CRC237 369799452/A11, CRC237 369799452/B21 to M.L.-K; CRC237 369799452/J to C.W., and as part of TRR 186 (Project Number 278001972) to H.E.), the German Federal Ministry of Education and Research (BMBF; 01GM2206C to M.L.-K.), and the Freie Universität Berlin (H.E.).

## Authors contribution

Conceptualization: H.M. and O.M.; Methodology, formal analysis and investigation: H.M., C.S., E.L.L., O.T., T.M., J.R., C.W., M.L.-K., D.M., F.H. and O.M.; Resources: D.M., H.E., C.K., M.L.-K., F.H., and O.M.; Data curation proteomics: D.M.; Writing - original draft: H.M., F.H. and O.M.; Writing - review and editing: H.M., E.L.L., O.T., C.W., D.M., M.L.- K., F.H., and O.M.; Supervision: F.H., and O.M.; Project administration: O.M.

## Supplementary information included

Materials request and correspondence scientific data: Olivia Majer

Correspondence biomedical data: Fabian Hauck

---

**Fig. S1:**
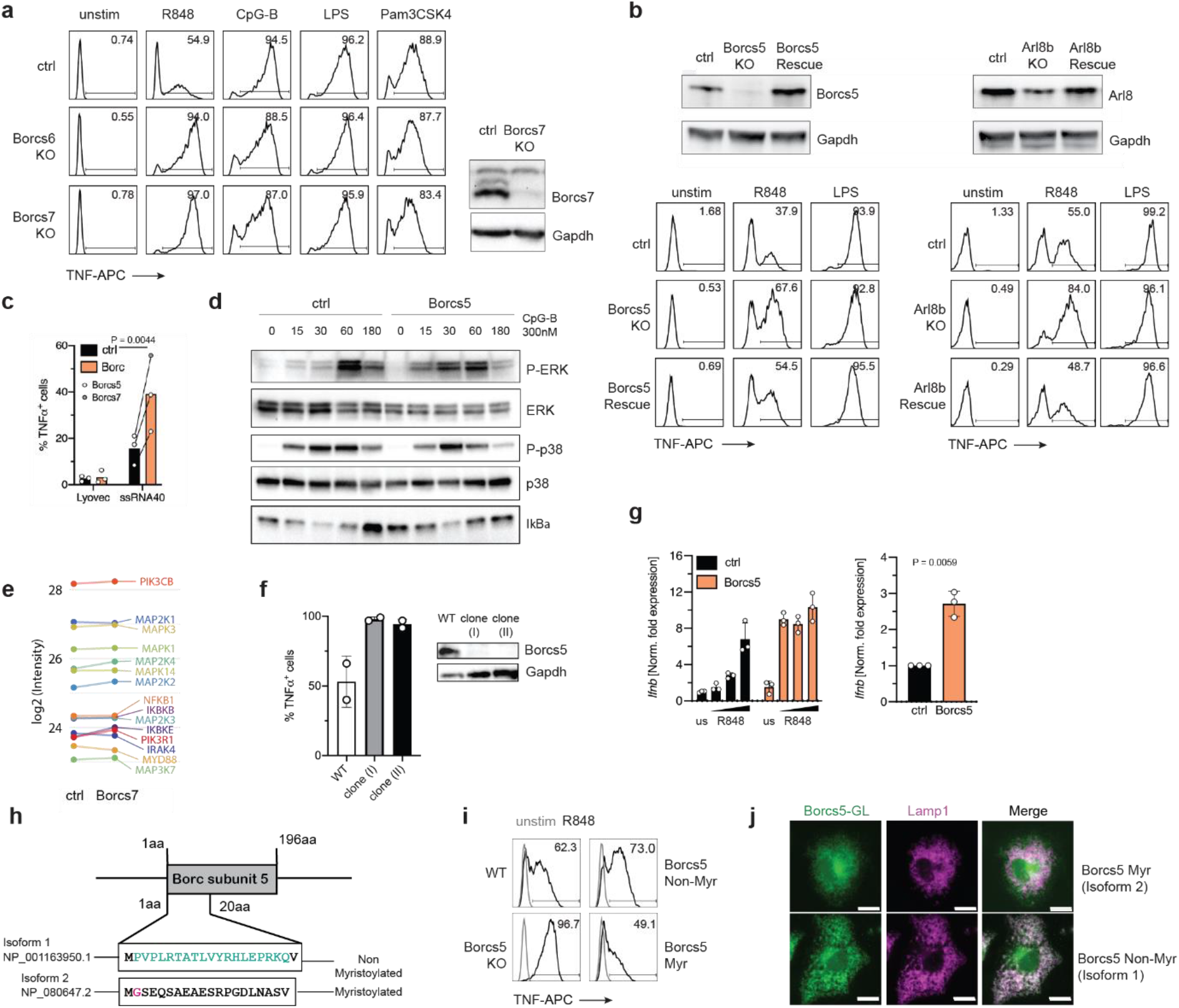
BORC deletion results in hyperactive TLR7 signaling. **a**, Intracellular TNF staining indicated Hoxb8 macrophage lines stimulated with R848 (10 ng/ml), CpG-B (300 nM), LPS (25ng/ml) or PamCSK4 (50ng/ml). Western blot shows validation of the Borcs7 knock-out. **b**, Intracellular TNF staining of Borcs5 (left panel) and Arl8b (right panel) rescue Hoxb8 macrophage lines stimulated with R848 (10 ng/ml) and LPS (25ng/ml). Western blots show re-expression of Borcs5 or Arl8b in the respective KO lines. Representative data of N=2. **c**, Intracellular TNF staining of indicated Hoxb8 macrophage lines with ssRNA40/LyoVec (5µg RNA/ml). Pooled data of N=3; ratio paired t test, two-tailed. **d**, Immunoblot of phosphorylated P-p38, P-ERK and IκBα of Hoxb8 macrophages stimulated with CpG-B for the indicated times. Representative data of N=2. **e**, Profile plots showing similar abundances (log2FC) of TLR signaling proteins between indicated Hoxb8 macrophage lines. Bars indicate SD of 4 replicate samples. **f**, Intracellular TNF staining of two independent Raw246.7 Borcs5 KO clones, stimulated with R848 (5 ng/ml). Western blot shows KO validation. **g**, Raw264.7 macrophage lines were stimulated with increasing concentrations of R848 (5, 10, 20g/ml) for 4h and *Ifnb* gene induction quantified by qPCR. Left graph: representative data. Right graph: pooled data for R848 10ng/ml of N=3; ratio paired t test, two-tailed. **h**, Schematic comparing N-terminal amino acid sequence of mouse Borcs5 isoform 1 and 2. **i**, Intracellular TNF staining of complemented Raw246.7 macrophage lines, stimulated with R848 (5 ng/ml). **j**, Representative images of Borcs5 KO Raw246.7 macrophages complemented with either Borcs5 isoform 1 or 2 and stained for GFP and Lamp1.

**Fig. S2:**
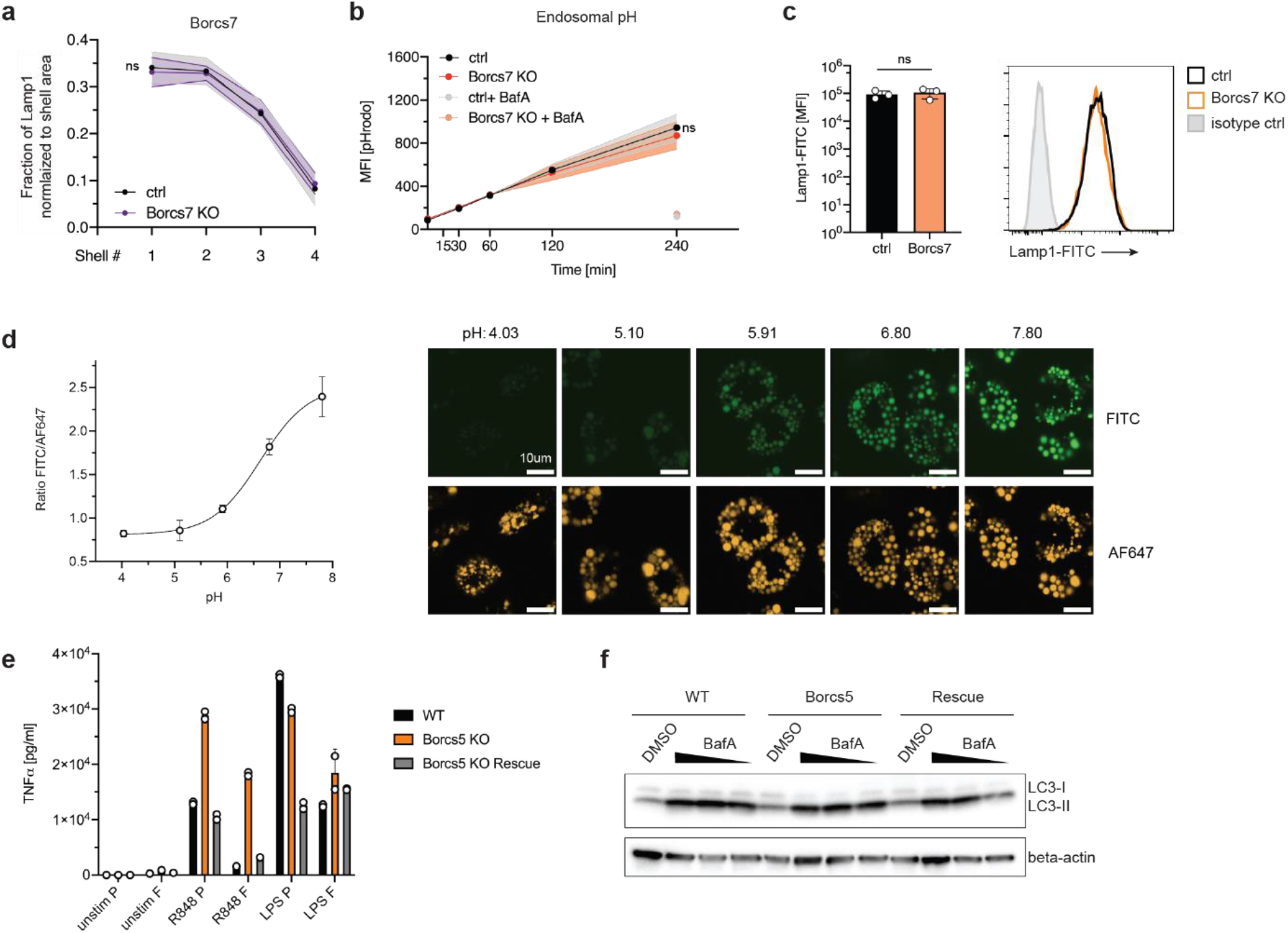
Similar endosome positioning, luminal pH and autophagic flux in BORC KO macrophages. **a**, Quantification of endosome radial distribution across segmented shells. Graphs show the fraction of late endosomes normalized per shell area. Pooled data from N=3, shaded areas indicate the SD. (Total number of cells analyzed: WT: 790, Borcs7 KO: 971); two-way ANOVA comparing variation between genotypes. **b**, Quantification of the luminal pH of endosomes by loading macrophages with the endosomal pH reporter pHrhodo Green (25µg/ml) for the indicated amounts of time. Bafilomycin A treatment (100nM) served as negative control. Pooled data of N=3. Shaded areas indicate the SD; two-way ANOVA comparing variation between genotypes. **c**, Lamp1 flow staining of Hoxb8 macrophages. Bar graph shows pooled data from N=3; unpaired t test, two-tailed. **d**, Representative calibration curve obtained by ratiometric fluorescence microscopy of Raw246.7 macrophages and representative images. **e**, TNF ELISA assay from cell culture supernatants of WT, Borcs5 KO and Borcs5 rescue Raw246.7 cells seeded on 10µg/ml fibronectin-coated(F) 6-well glass bottom plates or tissue culture plastic(P) treated with indicated amounts of ligands. Representative data of N=2. **f**, Autophagic flux was determined by incubating Raw macrophages of the indicated genotype with increasing amounts of Bafilomycin A (10, 50, 100nM) for 1h and reading out the accumulation of LC3-II by western. Representative image of N=2.

**Fig. S3:**
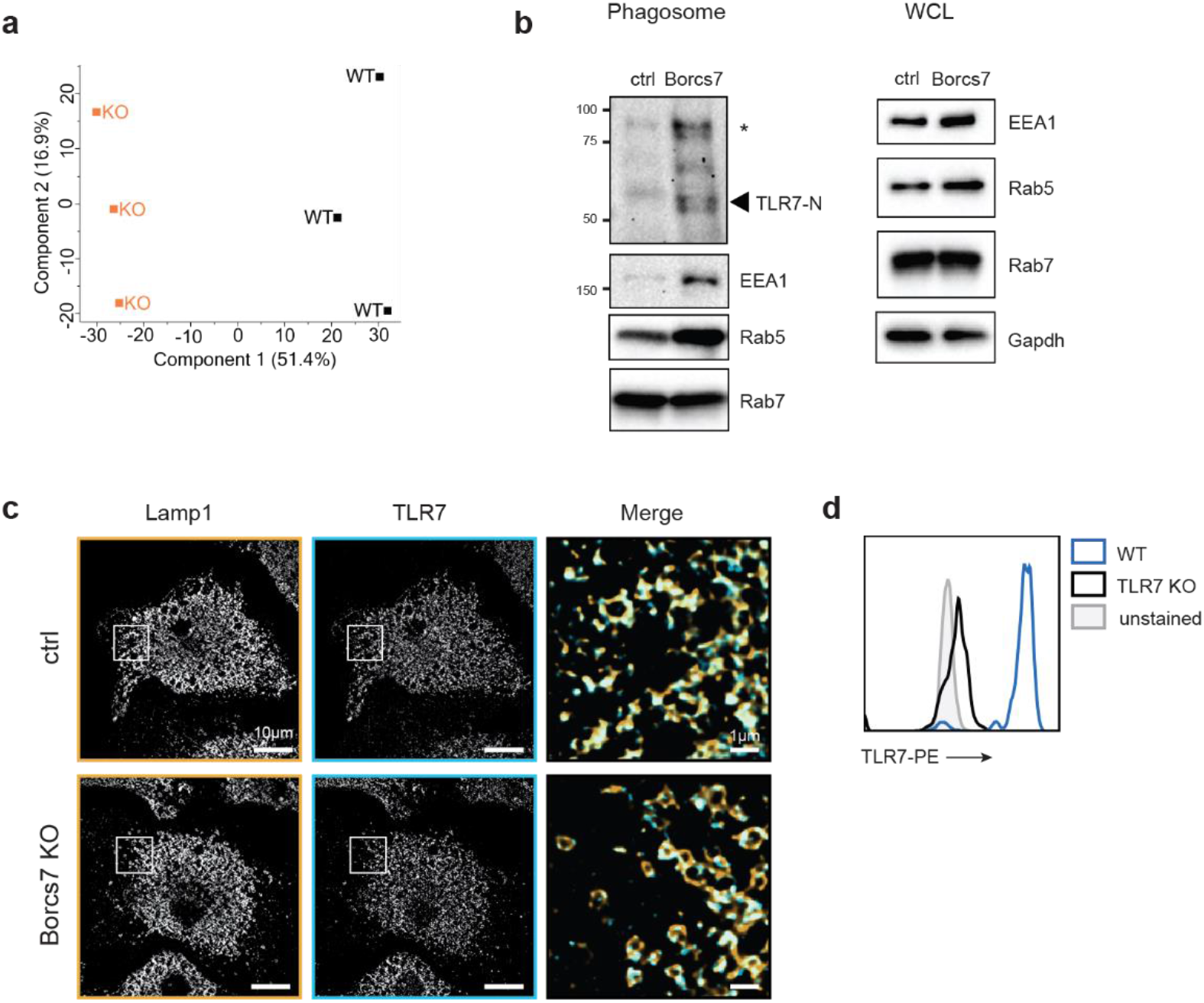
BORC-deficiency impairs endosome maturation and leads to TLR7 accumulation. **a**, Principal component analysis (PCA) plot of WT and Borcs5 phagosome analysis. **b**, Phagosome preparations form indicated Hoxb8 macrophage lines were analyzed by western blot. Representative data of N=2. WCL: whole cell lysate, FL: Full-length, C: cleaved. **c**, Representative images of indicated Hoxb8 macrophage lines stained for endogenous TLR7 and Lamp1 and imaged on a structured illumination microscope. **d**, Antibody specificity of anti-TLR7 was validated against TLR7 KO macrophages.

**Fig. S4:**
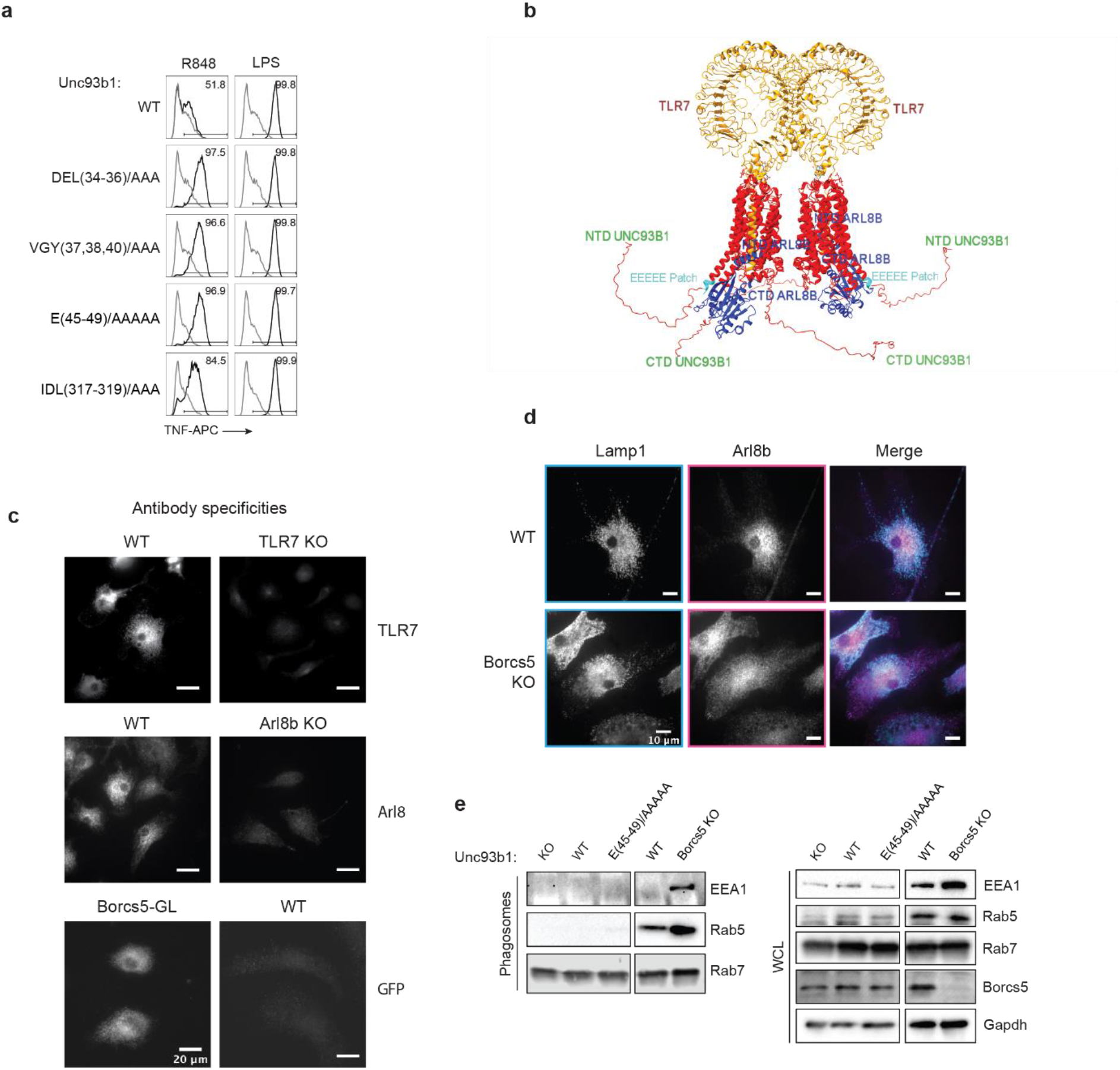
Mapping the Arl8b-Unc93b1 interaction. **a**, Intracellular TNF staining of Raw264.7 macrophage lines expressing the indicated Unc93b1 mutants stimulated with R848 (2ng/ml), CpG-B (300 nM), PolyIC (10µg/ml) or LPS (25ng/ml). **b**, AlphaFold multimer modeling of the TLR7-Unc93b1 complex and Arl8b. **c**, Antibody specificities of anti-TLR7, anti-Arl8b, and anti-GFP were validated against respective controls. Representative images with identical acquisition and display settings. **d**, Hoxb8 macrophages of the indicated genotypes were fixed and stained with anti-Lamp1 and anti-Arl8b. **e**, Western blot of phagosome preparations. Representative data of N=2. WCL: whole cell lysate.

**Fig. S5:**
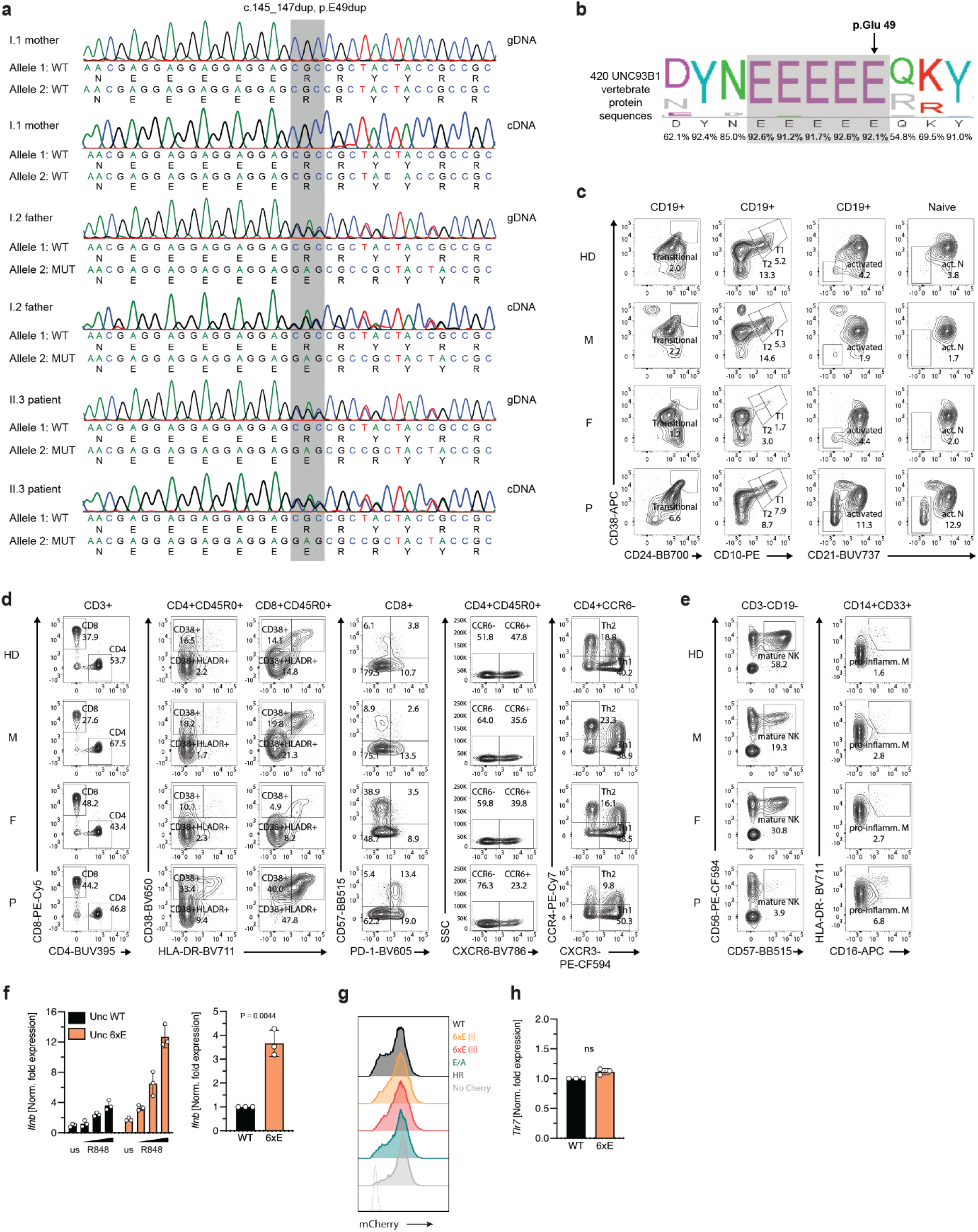
Clinical manifestation of the 6xE Unc93b1 mutation. **a,** Sanger sequencing electropherograms generated from PBMCs gDNA and cDNA. **b,** Consensus LogoPlot is depicted for the UNC93B1 5x glutamic acid stretch for 420 homologous vertebrate protein sequences containing 116 mammalian species. **c-e**, Flow cytometric immune phenotype of peripheral blood from a healthy donor (HD), the patient’s mother (M), father (F), and the patient (P). **f,** Raw264.7 macrophage lines were stimulated with increasing concentrations of R848 (5, 10, 20g/ml) for 4h and *Ifnb* gene induction quantified by qPCR. Left graph: representative data. Right graph: pooled data for R848 20ng/ml of N=3; ratio paired t test, two-tailed. **g**, Equal expression levels of Unc93b1 variants monitored through an IRES-mCherry. **h**, qPCR of *Tlr7* in unstimulated Raw264.7 macrophage lines. Pooled data of N=3; ratio paired t test, two-tailed.

**Fig. S6:**
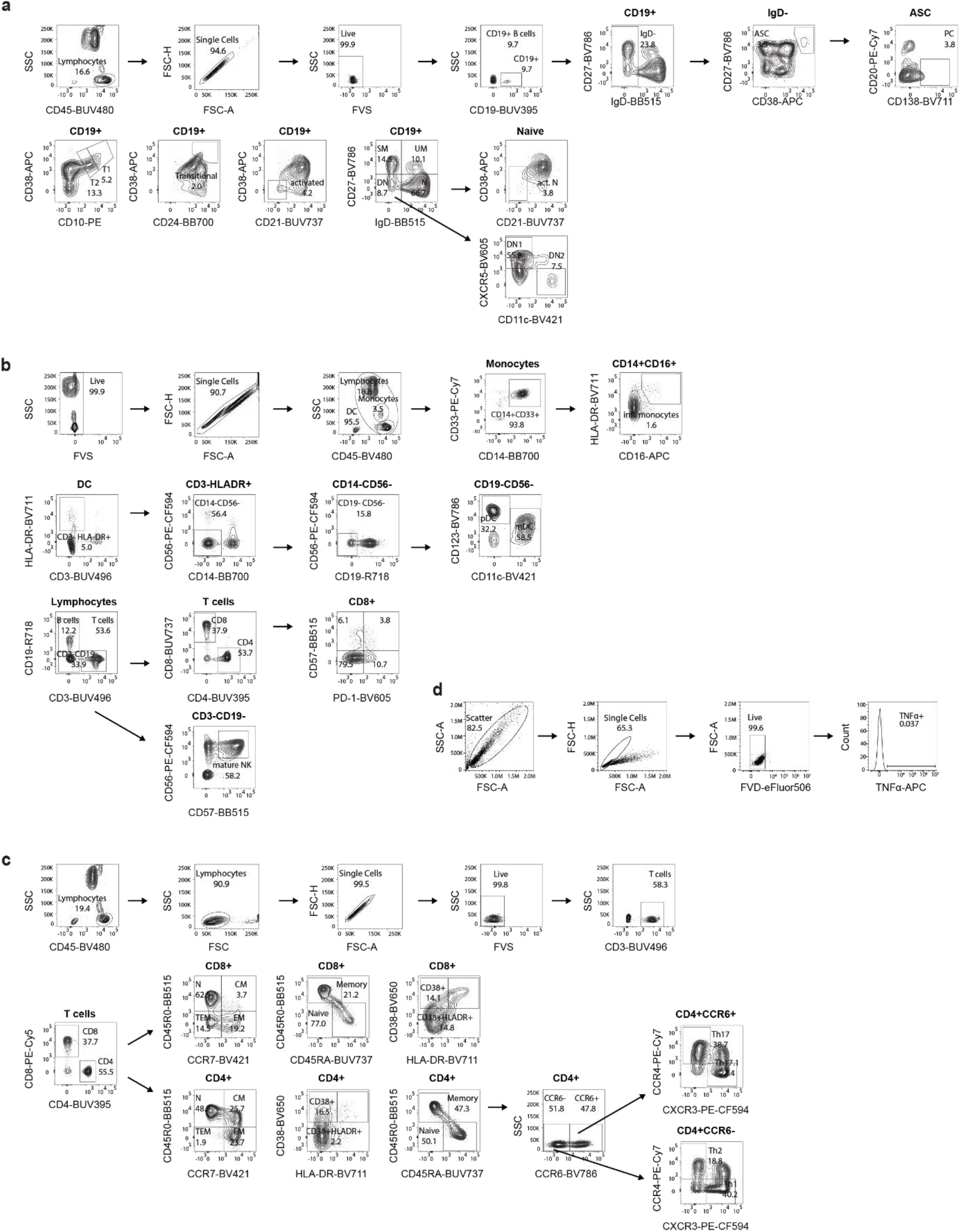
Gating strategies. Gating strategies used in **a**, Fig. 5e and Fig. S5c for B-cell subsets. **b**, Fig. 5f and Fig. S5e for monocytes, dendritic cells, NK cells and T cell subsets. **c**, Fig. S5d for T cell subsets. **d**, General gating strategy for TNF intracellular cytokine staining in mouse macrophages. Same gating also applies to intracellular TLR7-PE, Lamp1-FITC staining.

