## Supplementary Information for "Endosome dysfunction leads to gain-of-function TLR7 and human lupus"

Includes:

**Extended case report**

**Supplementary data 1-2**

**Supplementary methods**

1. Exome sequencing
2. Sanger sequencing and RT-PCR
3. Phylogenetic analysis
4. Immunoblotting of PBMCs
5. Quantitative real-time PCR of interferon-stimulated genes
6. Analysis endosome positioning
7. Analysis ratiometric fluorescence microscopy
8. Mass Spectrometry:
   1. Proteomics Sample Preparation with Label-Free Quantification (LFQ)
   2. Mass Spectrometry Settings for Proteome Profiling with the Q Exactve HF
   3. Mass Spectrometry Settings for Proteome Profiling with the timsTOF SCP
   4. Experimental Design, Statistical Rationale, Pathway, and Data Analyses

**Supplementary tables**

1. Laboratory data of P1 (separate file)
2. Proteome analysis of ctrl and Borcs7 KO Hoxb8 macrophages: MaxQuant processed files (separate file)
3. Key resources table

**Source data**

Western blots – original images (separate file)

**Extended case report**

An 11-year-old female patient (P1) from a non-consanguineous Caucasian family was referred for suspected systemic lupus erythematosus (SLE) based on the EULAR/ACR 2019 SLE classification criteria^1^. She had a medical antecedent of atopic dermatitis (onset at 6 years of age), Hashimoto’s thyroiditis (onset at 7 years of age, Suppl. Tab.1 and Suppl. Data 1a), generalized lymphadenopathy (cervical, axillary, inguinal, abdominal, retroperitoneal, onset at 10 years of age, Suppl. Data 1c) and sialadenitis (parotid and submandibular glands, onset at 10 years of age, Suppl. Data 1b). At evaluation, she was found to have antinuclear antibodies (ANA > 1:12,800) as SLE diagnostic entry criterion, and additive SLE diagnostic criteria such as leucopenia (3.63 G/l, 3 points), proteinuria (225 mg/m^2^, 4 points), low C3 and C4 (C3 0.52 g/l, C4 <0.02 g/l, 4 points), and anti-ds-DNA antibodies (461 IE/ml, 6 points) (Suppl. Tab.1) that summed up to 17 points thereby exceeding the required ≥10 points for a definitive SLE diagnosis^1^. In addition, she had splenomegaly (Suppl. Data 1d), massive hypergammaglobulinemia (IgM 4.99 g/l and IgG 64.5 g/l), anti-C1q- (22 U/ml), anti-SSA/Ro (positive) and SSB/La- (positive antibodies (Suppl. Tab.1). To identify the genetic cause of P1’s SLE, we performed karyotyping (data not shown), array-CGH (data not shown) and trio exome sequencing (data not shown). We identified no structural and no genetic variants in genes known to be associated with SLE or diseases of immune dysregulation^2^. However, we detected a monoallelic UNC93B1 (ENST00000227471) c.145_147dup p.Glu49dup variant of unknown significance (VUS, ACMG standards and guidelines^3^) in P1 and her father who had not yet come to medical attention (Fig. 5a-b). UNC93B1 p.Glu49dup was neither present in public databases (GnomAD, ExAC and GME) nor in or our in-house exome database (5,600 exomes) and thus was a private variant with a minor allele frequency (MAF) <10^-5^. We confirmed UNC93B1 p.Glu49dup in P1 and her father by Sanger sequencing of genomic DNA and tested its expression by reverse transcription of total PBMC mRNA and Sanger sequencing of cDNA (Fig. S5a). UNC93B1 p.Glu49dup extends an N-terminal 5x Glu stretch (UniProtKB Q9H1C4, amino acids 45-49) preceding (Glu 45-46) or taking part of the first transmembrane α-helix into (Glu 47-49) into a 6x Glu stretch^4^ (Fig. 5a-b). Phylogenetic analysis of 420 homologous UNC93B1 vertebrate protein sequences encompassing 116 mammalian sequences showed high conservation of the 5x Glu stretch (Fig. 5c and Fig. S5b). To probe if the monoallelic UNC93B1 variant alters protein expression, we isolated whole protein from primary PBMC and performed western blotting. UNC93B1 protein levels were comparable for P1 and her parents (Fig. 5d). As UNC93B1 regulates TLR7-signaling to prevent SLE-like autoimmunity^5-7^, we measured type 1 interferon-related biomarkers with real-time quantitative PCR (RT-qPCR)^8^ in samples from P1 and her parents. While the mother displayed normal values, P1 and her father had an increased interferon signature (Fig. 5g) segregating with the monoallelic UNC93B1 p.Glu49dup. Deep immune phenotyping of PBMCs showed accumulation of total CD19^+^ B-cells, IgD^-^CD27^-^ double-negative (DN) B-cells and their CXCR5^-^CD11c^+^ DN2 subset [i.e., age-associated B-cells (ABC)], and CD20^-^CD138^+^ plasma cells in the patient and to some extend in her father being compatible with increased TLR7 signaling (Fig. 5e)^9^. CD24^+^CD38^+^ transitional B-cells and their CD10^+^CD38^+^ transitional 1 subset, total CD19^+^ B- and naïve CD21^low^CD38^low^ activated B-cells were increased in the patient (Fig. S5c). In addition, CD4^+^/CD8^+^ T-cell ratio, CD4^+^ and CD8^+^CD38^+^HLA-DR^+^ activated T-cells, and CD8^+^CD57^+^PD-1^+^ senescent T-cells were increased in the patient and to some extend in her father reflecting reactive T-cell states. In the patient, CD4^+^CD45R0^+^CCCR6^-^ helper T-cells and their CCR4^+^CXCR3^-^ helper type 2 (Th2) subset was decreased reflecting lupus disease activity (Fig. S5d)^10,11^. CD16^+^CD56^+^ NK-cells and CD56^+^CD57+ mature NK-cells were reduced, and CD14^+^CD16^-^ classical monocytes were reduced and CD16^+^HLA-DR^+^ pro-inflammatory monocytes were increased in the patient (Fig. 5f), while lin^-^CD123^+^ plasmacytoid dendritic cells (pDCs) were reduced in the patient and her father reflecting an inflammatory state (Fig. 5f)^10,12^.

**Supplementary Data 1**

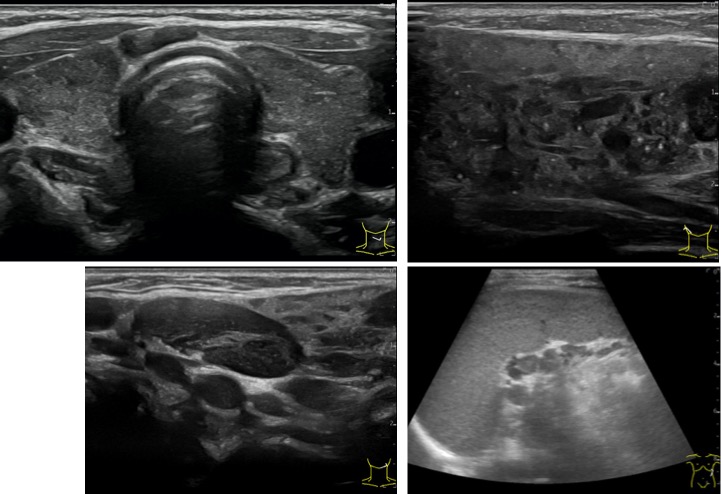

**d**

**c**

**b**

**a**

**Radiological correlates of multiorgan autoimmunity and generalized lymphoproliferation.**

**a,** Inhomogeneous parenchyma of the thyroid gland compatible with Hashimoto’s thyroiditis. **b,** Inhomogeneous parenchyma of the submandibular gland with multiple calcifications compatible with chronic sialadenitis. **c,** Increased number and size (max. 22.0 cm x 0.9 cm) of cervical lymph nodes representing generalized lymphadenopathy. **d,** Splenomegaly (11.2 cm length) with inhomogeneous, fine grained parenchyma and increased number and size (max. 1.0 cm x 0.8 cm) of hilar lymph nodes reflecting lymphoproliferation.

**Supplementary Data 2**

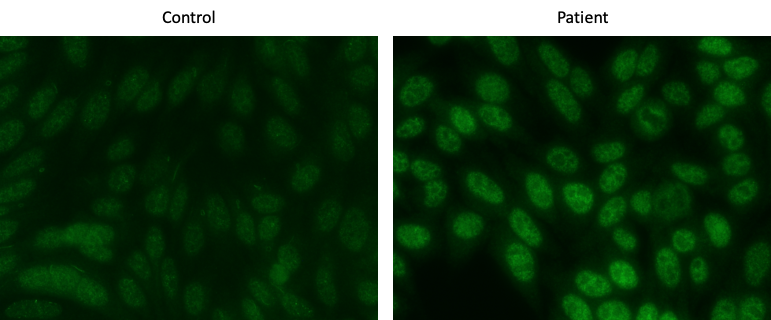

Immunofluorescences analysis of control and patient serum on HEp-2 cells (630x magnification) showing a negative staining and a speckled pattern of antinuclear antibodies.

**Supplementary methods**

**Exome sequencing**

Exome sequencing for was performed at the Dr. von Hauner Children's hospital next-generation-sequencing facility. Briefly, genomic DNA from whole blood was used for preparation of whole-exome libraries using the SureSelect XT Human All Exon V6+UTR kit (Agilent Technologies,) and subsequently sequenced with a NextSeq 500 platform (Illumina) to an average coverage depth of 90x. Bioinformatic analysis used Burrows-Wheeler Aligner (BWA 0.7.15), Genome Analysis ToolKit (GATK 3.6) and Variant Effect Predictor (VEP 89). For allele frequency filtering we used public (e.g., GnomAD and ExAC) and in-house databases (approx. 5,600 exomes).

**Sanger sequencing and RT-PCR**

Genomic DNA was isolated from whole blood samples with the iPrep PureLink gDNA Blood kit (Life Technologies) and the UNC93B1 variant was confirmed by Sanger sequencing (Eurofins, Ebersberg, Germany) of genomic DNA from P1 and available family members (forward primer: 5´-GCAGGGCGACGAGGACCTGCTC-3´; reverse primer: 5´-CGCCGGCCTCCAATTCTGACGGTG-3´). Total mRNA was isolated from PBMCs using Qiazol lysis reagent (Qiagen, Hilden, Germany), reverse transcribed into cDNA (QuantiTect, Qiagen) and the UNC93B1 variant was confirmed by Sanger sequencing (Eurofins) of cDNA from P1 and available family members (forward primer: 5´-GCAGGGCGACGAGGACCTGCTC-3´; reverse primer: 5´-GCGATGGGAGTCACGTTGATG-3´).

**Phylogenetic analysis**

500 homologous protein sequences for human UNC93B1 (Q9H1C4) were identified by blast search in Uniprot^13^ and imported to Jalview 2.11^14^. Hereupon, 420 unique protein sequences were identified by selecting full-length or canonical sequences manually in case of intraspecies redundancies and aligned by ClustalOWS^15^ in Jalview 2.11. The LogoPlot diagram was constructed with Jalview 2.11 and Inkscape 1.2 (Inkscape Project, 2020. *Inkscape*, Available at: https://inkscape.org).

**Immunoblotting of PBMCs**

Whole cell lysates were prepared from PBMCs. Equal protein amounts were separated by SDS-PAGE, transferred to nitrocellulose and probed with anti-human-UNC93B1 (polyclonal, 1:1000, PA5-20510, ThermoFisher Scientific, Waltham Massachusetts, USA), anti-human-GAPDH (6C5, 1:3000, sc-32233, Santa Cruz, Dallas, USA) and HRP-conjugated anti-rabbit (sc-2357, 1:10000) and anti-mouse secondary antibodies (sc-2005, 1:10000, both Santa Cruz).

**Quantitative real-time PCR of interferon-stimulated genes**

Interferon-signature was done as previously described^8^.

**Analysis endosome positioning**

To quantify and analyse the distribution of lysosomes within the cells, a specific pipeline in CellProfiler^16^ was created. The designed pipeline divides the cytoplasm of each cell into four automatically defined shells and then quantifies the Lamp1 signal within each shell. The pipeline included the following steps: First, the nuclei of cells (the center points are used to generate the shells) as well as the cell boundaries were segmented. To exclude dividing cells or other outliers, a manual editing step was included in the pipeline. Next, the build-in module “RadialDistribution_MeanFrac” was used to automatically generate the shells and quantify the lysosomal distribution between them. The “RadialDistribution_MeanFrac” is automatically calculated by CellProfiler as fraction of total intensity normalized by fraction of pixels at a given radius.

**Analysis ratiometric fluorescence microscopy**

For each experiment, a pH calibration curve was generated before acquiring experimental data. The calibration buffers contained 143mM KCl, 5mM glucose, 1mM MgCl_2_ and 20mM HEPES. The buffer solutions were adjusted to the final pH ranging between 4.0 and 8.0 by using 1 M NaOH or 1 M HCl. All buffers were filtered before use. Nigericin (10µM, N7143, Sigma) and Monensin (5µM, M5273, Sigma) were added to the buffers right before use. To record the calibration curve, cells were Dextran-loaded and incubated in CellMask as described above. After a final PBS wash, cells were incubated in the respective pH calibration buffer for 10mins and then imaged immediately. Calibration was performed sequentially on the same cells starting from the most alkaline to the most acidic buffer.

To quantify the pH of endosomes, a custom-made Cell Profiler pipeline was developed: First, the images were background subtracted using a dark frame image. Then, a median filter was applied on the CellMask channel and inverted for segmentation of cells. The FITC- and 647-labelled endosomes inside the cells were segmented as objects, shrunken to a single pixel, expanded by 1 pixel on all sides and related to each other. The segmented and related endosomes were expanded by 2 more pixels on all sides before measuring the intensities in each channel. The ratio of fluorescence intensities of both channels was calculated. All endosomes belonging to the same cell were averaged to express the mean endosomal pH per single cell. For calibration the measured average ratios of endosomes (Y) incubated in known pH (X) buffers were plotted to generate a standard curve. The unknown pH of cells was interpolated using the calculated ratios.

**Mass Spectrometry**

**Proteomics Sample Preparation with Label-Free Quantification (LFQ)**

Proteomics sample preparation was done according to a published protocol with minor modifications^17^. Eight biological replicates of Borcs7^-/-^ and vector control cells (2 million per sample) were lysed under denaturing conditions in a buffer containing 3 M guanidinium chloride (GdmCl), 10 mM tris(2-carboxyethyl)phosphine (TCEP), 40 mM chloroacetamide and 100 mM Tris-HCl pH 8.5. Lysates were denatured at 95°C for 10 min shaking at 1000 rpm in a thermal shaker and sonicated in a water bath for 10 min. Each lysate was diluted with a dilution buffer containing 10% acetonitrile and 25 mM Tris-HCl, pH 8.0, to reach a 1 M GdmCl concentration. Then, proteins were digested with LysC (Roche, Basel, Switzerland; enzyme to protein ratio 1:50, MS-grade) shaking at 700 rpm at 37°C for 2.5 hours. The digestion mixture was diluted again with the same dilution buffer to reach 0.5 M GdmCl, followed by a tryptic digestion (Roche, enzyme to protein ratio 1:50, MS-grade) and incubation at 37°C overnight in a thermal shaker at 700 rpm. Peptide desalting was performed according to the manufacturer’s instructions (Pierce C18 Tips, Thermo Scientific, Waltham, MA). Desalted peptides were reconstituted in 0.1% formic acid in water and further separated into four fractions by strong cation exchange chromatography (SCX, 3M Purification, Meriden, CT). Eluates were first dried in a SpeedVac, then dissolved in 5% acetonitrile and 2% formic acid in water, briefly vortexed, and sonicated in a water bath for 30 seconds before injection to nano-LC-MS.

**Mass Spectrometry Settings for Proteome Profiling with the Q Exactve HF**

LC-MS/MS was carried out by nanoflow reverse phase liquid chromatography (Dionex Ultimate 3000, Thermo Scientific) coupled online to a Q-Exactive HF Orbitrap mass spectrometer (Thermo Scientific), as reported previously^18^. Briefly, the LC separation was performed using a PicoFrit analytical column (75 μm ID × 50 cm long, 15 µm Tip ID; New Objectives, Woburn, MA) in-house packed with 3-µm C18 resin (Reprosil-AQ Pur, Dr. Maisch, Ammerbuch, Germany). Peptides were eluted using a gradient from 3.8 to 38% solvent B in solvent A over 120 min at 266 nL per minute flow rate. Solvent A was 0.1 % formic acid and solvent B was 79.9% acetonitrile, 20% H_2_O, and 0.1% formic acid. For the IP samples, a one-hour gradient was used. Nanoelectrospray was generated by applying 3.5 kV. A cycle of one full Fourier transformation scan mass spectrum (300−1750 m/z, resolution of 60,000 at m/z 200, automatic gain control (AGC) target 1 × 10^6^) was followed by 12 data-dependent MS/MS scans (resolution of 30,000, AGC target 5 × 10^5^) with a normalized collision energy of 25 eV. To avoid repeated sequencing of the same peptides, a dynamic exclusion window of 30 sec was used. In addition, only peptide charge states between two to eight were sequenced.

Raw MS data were processed with MaxQuant software (v2.0.1.0) and searched against the mouse proteome database UniProtKB with 55,366 entries, released in March 2021. Parameters of MaxQuant database searching were a false discovery rate (FDR) of 0.01 for proteins and peptides, a minimum peptide length of seven amino acids, a first search mass tolerance for peptides of 20 ppm and a main search tolerance of 4.5 ppm, and using the function “match between runs”. A maximum of two missed cleavages was allowed for the tryptic digest. Cysteine carbamidomethylation was set as fixed modification, while N-terminal acetylation and methionine oxidation were set as variable modifications. Contaminants, as well as proteins identified by site modification and proteins derived from the reversed part of the decoy database, were strictly excluded from further analysis. The MaxQuant processed output files can be found in Supplementary table 2: Proteome analysis of ctrl and Borcs7 Hoxb8 macrophages, showing peptide and protein identification, accession numbers, % sequence coverage of the protein, q-values, and LFQ intensities. The mass spectrometry data have been deposited to the ProteomeXchange Consortium (http://proteomecentral.proteomexchange.org) via the PRIDE partner repository^19^ with the dataset identifier PXD039263.

**Mass Spectrometry Settings for Proteome Profiling with the timsTOF SCP**

Phagosome preparations from WT and Borcs5 KO Raw264.7 macrophages were prepared in triplicates as described in Materials and Methods “Phagosome preparation”. Instead of denaturing proteins off the beads, bead-containing intact phagosomes were pelleted, washed 4x in cold 100 mM ammonium bicarbonate (NH₄HCO₃), and snap-frozen.

100 µl of fresh lysis buffer was added and digested as mentioned in the section Proteomics Sample Preparation with LFQ. Peptide desalting was performed on self-prepared C18 Tips. Samples were acidified (final formic acid concentration of 2 %), shortly vortexed, and centrifuged. The tubes were placed on a magnetic rack and the supernatant was loaded on the activated and equilibrated C18 columns. The flow through was loaded again once, the column was washed with 0.1% formic acid and peptides were eluted with 60% acetonitrile and 0.1% formic acid in water. 10% of eluates were first dried in a SpeedVac and, then loaded onto Evotips Pure (Evosep, Odense, Denmark) tips according to the manufacturer’s protocol. Peptide separation was carried out by nanoflow reverse phase liquid chromatography (Evosep One, Evosep) using the Endurance column (15 cm x 150 µm ID, with Reprosil-Pur C18 1.9 µm beads #EV1106, Evosep) with the 30 samples a day method (30SPD). The LC system was online coupled to a timsTOF SCP mass spectrometer (Bruker Daltonics, Bremen, Germany) applying the data-independent acquisition (DIA) with parallel accumulation serial fragmentation (PASEF) method. MS data were processed with Dia-NN (v1.8) and searched against an *in silico* predicted mouse spectra library. The “match between run” feature was used. A t-test with Benjamini-Hochberg correction was performed by Perseus (v1.6.15.0) on normalized protein values to identify significant regulations between KO and controls.

The mass spectrometry data have been deposited to the ProteomeXchange Consortium (http://proteomecentral.proteomexchange.org) via the PRIDE partner repository^19^ with the dataset identifier PXD040042.

**Experimental Design, Statistical Rationale, Pathway, and Data Analyses**

The correlation analysis of biological replicates and the calculation of significantly different metabolites and proteins were done with Perseus (v1.6.15.0). LFQ intensities, originating from at least two different peptides per protein group (e.g. KO versus control), were transformed by log_2_. Only groups with valid values in at least one group were used, missing values were replaced by values from the normal distribution. Statistical analysis was done by a two-sample t-test with Benjamini-Hochberg (BH, FDR of 0.05) correction for multiple testing. Significantly regulated proteins between Hoxb8 Borcs7 KO and vector controls were indicated by a plus sign in Supplementary Table 2.

A gene set enrichment analysis (GSEA, v4.1.0)^20^ was applied to see, if *a priori* defined sets of proteins show statistically significant, concordant differences between Borcs7 and controls. All proteins with ratios calculated by Perseus were used for GSEA analysis. GSEA standard settings were used, except that the minimum size exclusion was set to 5, and Reactome and KEGG v7.4 were used as gene set databases. The cutoff for significantly regulated pathways was set to be ≤ 0.05 p-value and ≤ 0.05 FDR.

**Table S1. Laboratory data of P1 (separate file)**

**Table S2. Proteome analysis of ctrl and Borcs7 KO Hoxb8 macrophages: MaxQuant processed files (separate file)**

Table S3. Key resources table

| **Key resources table** | | |
| --- | --- | --- |
| **Plasmids** | | |
| **Insert** | **Backbone** | **Marker** |
| Borcs5 mGreenlantern Myristoylated | MSCV (Takara) | mGreenlantern |
| Borcs5 - mGreenLantern non-myristoylated | MSCV (Takara) | mGreenlantern |
| Arl8b rescue* | MSCV (Takara) | Thy1.1 |
| Arl8b overexpression | pQCXI | Puromycin |
| Borcs5 rescue* Myristoylated | MSCV (Takara) | Thy1.1 |
| gRNA | lentiGuide-puro (Addgene #52963) |  |
| VSV-G | pCMV (Addgene #8454) | Ampicillin |
| psPAX2 | psPAX2 (Addgene #12260) | Ampicillin |
| Mouse TLR3-HA (codon-optimized) | MSCV (Takara) | Thy1.1 |
| Mouse TLR7-HA (codon-optimized) | MSCV (Takara) | Thy1.1 |
| Mouse TLR9-HA | MSCV (Takara) | Thy1.1 |
| Mouse Unc93b1 WT (codon-optimized) | MSCV (Takara) | mCherry-T2A-Puromycin |
| Mouse Unc93b1 H412R | MSCV (Takara) | mCherry-T2A-Puromycin |
| Mouse Unc93b1 DEL(34-36)/AAA | MSCV (Takara) | mCherry-T2A-Puromycin |
| Mouse Unc93b1 VGY(37,38,40)/AAA | MSCV (Takara) | mCherry-T2A-Puromycin |
| Mouse Unc93b1 E(45-49)/AAAAA | MSCV (Takara) | mCherry-T2A-Puromycin |
| Mouse Unc93b1 IDL(317-319)/AAA | MSCV (Takara) | mCherry-T2A-Puromycin |
| Mouse Unc93b1 6xE (Glu49dup) | MSCV (Takara) | mCherry-T2A-Puromycin |
| **gRNAs** | | |
| Mouse Borcs5 guide 1 | GGCCAAGATGGACGATATCG | |
| Mouse Borcs5 guide 2 | GTCATTGCTGACATTCCGTG | |
| Mouse Arl8b | CACCGCGATGACATTGACGA | |
| Mouse Borcs6 | TCGACGGCAGACCCTCAGAG | |
| Mouse Borcs7 | AGCAGGTGCTGAAAGGCTCG | |
| **qPCR primers (All from PrimerBank)** | | |
| **Mouse:** | | |
| mIFNbeta F | AGCTCCAAGAAAGGACGAACA | |
| mIFNbeta R | GCCCTGTAGGTGAGGTTGAT | |
| mGapdh F | AGGTCGGTGTGAACGGATTTG | |
| mGapdh R | GGGGTCGTTGATGGCAACA | |
| mHPRT F | TCAGTCAACGGGGGACATAAA | |
| mHPRT R | GGGGCTGTACTGCTTAACCAG | |
| mTLR7 F | ATGTGGACACGGAAGAGACAA | |
| mTLR7 R | ACCATCGAAACCCAAAGACTC | |
| mBorcs7 F | TCGCAAGCCCGATTTGGTC | |
| mBorcs7 R | CTTGGTGAGCGCGATTACAT | |
| mVps41 F | CCCAAACTGAAGTATGAAAGGCT | |
| mVps41 R | TTCCCAGTGCCAAAAACTTGT | |
| mVps39 F | TCTGCGTTGGTTTCAAGAGAG | |
| mVps39 R | GGGCAACTAAGGGCTCCAG | |
| **Human:** | | |
| hIL8 F | ACTGAGAGTGATTGAGAGTGGAC | |
| hIL8 R | AACCCTCTGCACCCAGTTTTC | |
| hGAPDH F | ACAACTTTGGTATCGTGGAAGG | |
| hGAPDH R | GCCATCACGCCACAGTTTC | |
| hHRPT F | CCTGGCGTCGTGATTAGTGAT | |
| hHRPT R | AGACGTTCAGTCCTGTCCATAA | |
| **SDM primer (designed with the online NEBaseChanger tool)** | | |
| Unc93b1 6xE F | gaaAGGCGCTACTACAGAAGA | |
| Unc93b1 6xE R | CTCTTCCTCTTCCTCGTTG | |
| **siRNA (all from Dharmacon, siGenome siRNA SMARTPool)** | | |
| human Borcs5 | Entrez Gene 118426 | |
| mouse Borcs7 | Entrez Gene 66439 | |
| mouse Vps39 | Entrez Gene 269338 | |
| mouse Vps41 | Entrez Gene 218035 | |
| Non-Targeting Control siRNA | #3 | |

* for gene rescue, silent mutations were included within the PAM site and adjacent
