## Supplementary material for "Endosome dysfunction leads to gain-of-function TLR7 and human lupus": Source data

### Slide 1
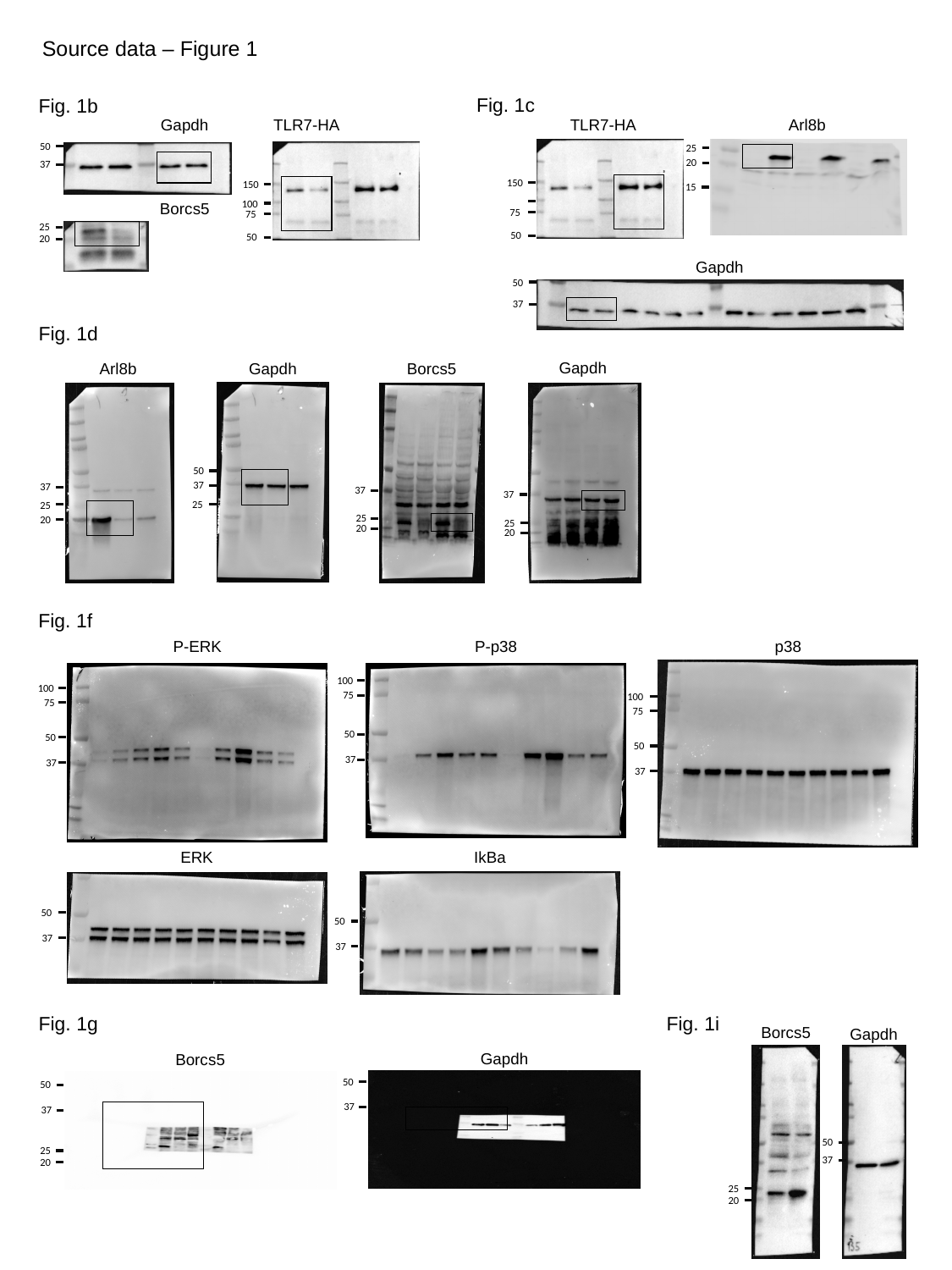

Source data – Figure 1
Fig. 1c
Fig. 1b
Gapdh
TLR7-HA
TLR7-HA
Arl8b
50
25
20
37
150
150
15
100
100
Borcs5
75
75
25
50
50
20
Gapdh
50
37
Fig. 1d
Gapdh
Arl8b
Gapdh
Borcs5
50
37
37
37
37
25
25
25
20
25
20
20
Fig. 1f
P-ERK
P-p38
p38
100
100
75
100
75
75
50
50
50
37
37
37
ERK
IkBa
50
50
37
37
Fig. 1g
Fig. 1i
Borcs5
Gapdh
50
37
25
20
Gapdh
Borcs5
50
50
37
37
25
20

### Slide 2
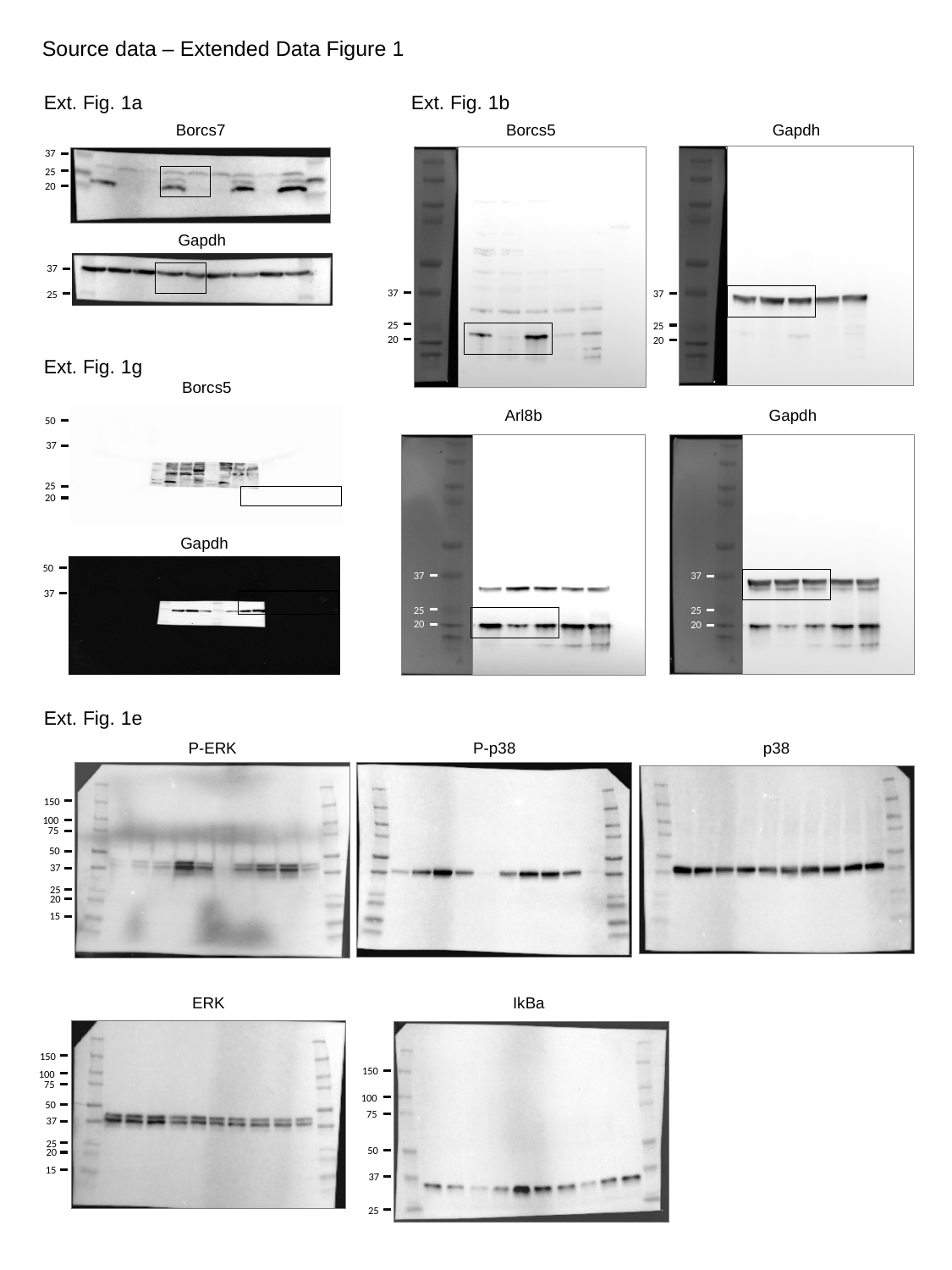

Source data – Extended Data Figure 1
Ext. Fig. 1a
Ext. Fig. 1b
Borcs7
Borcs5
Gapdh
37
25
20
Gapdh
37
37
37
25
25
25
20
20
Ext. Fig. 1g
Borcs5
Arl8b
Gapdh
50
37
25
20
Gapdh
50
37
37
37
25
25
20
20
Ext. Fig. 1e
P-ERK
P-p38
p38
150
100
75
50
37
25
20
15
ERK
IkBa
150
150
100
75
100
50
75
37
25
50
20
15
37
25

### Slide 3
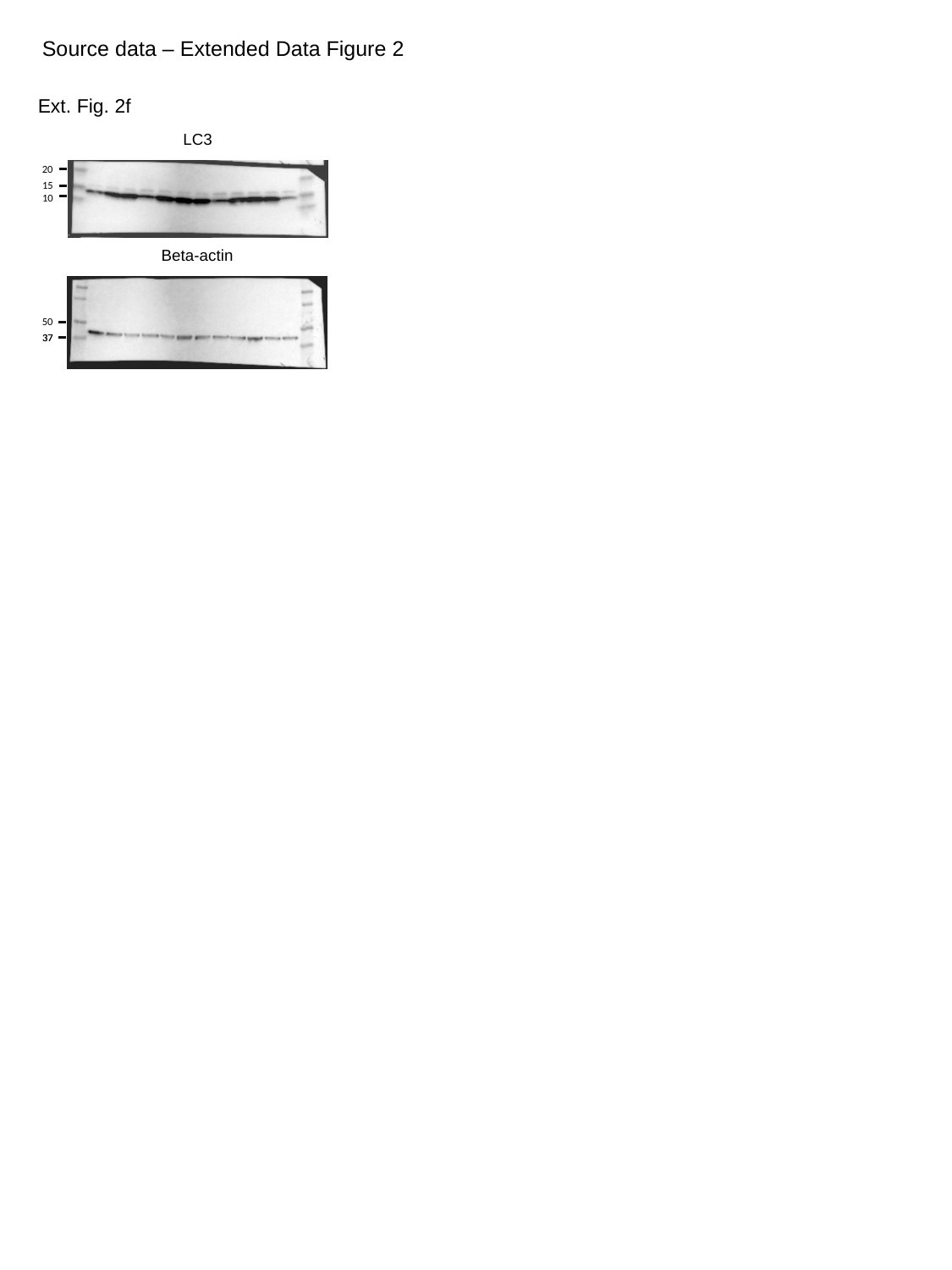

Source data – Extended Data Figure 2
Ext. Fig. 2f
LC3
20
15
10
Beta-actin
50
37
37

### Slide 4
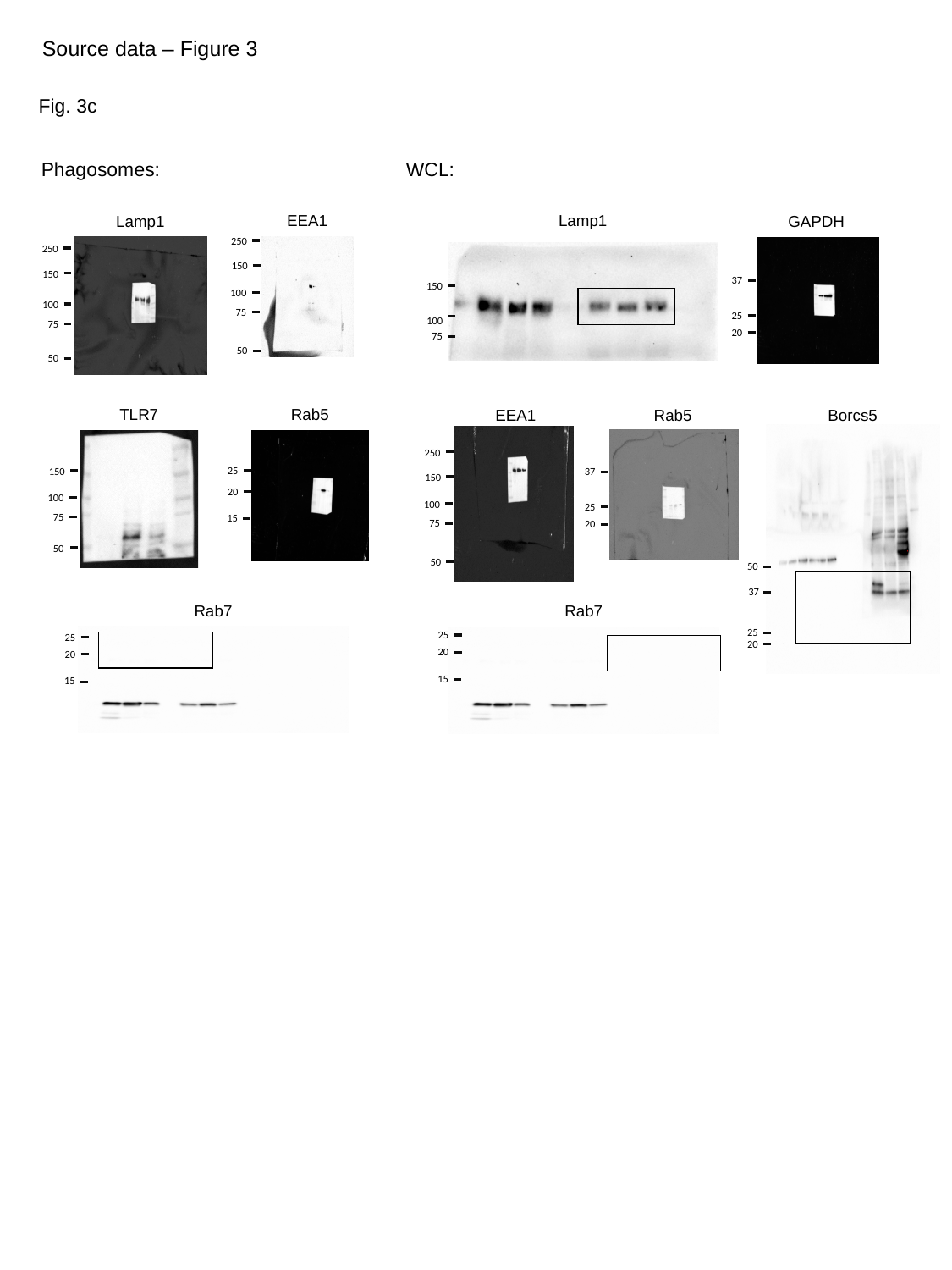

Source data – Figure 3
Fig. 3c
WCL:
Phagosomes:
EEA1
Lamp1
Lamp1
GAPDH
250
250
150
150
37
150
100
100
75
25
100
75
20
75
50
50
TLR7
Rab5
Borcs5
EEA1
Rab5
250
25
150
37
150
20
100
100
25
75
15
75
20
50
50
50
37
Rab7
Rab7
25
25
25
20
20
20
15
15

### Slide 5
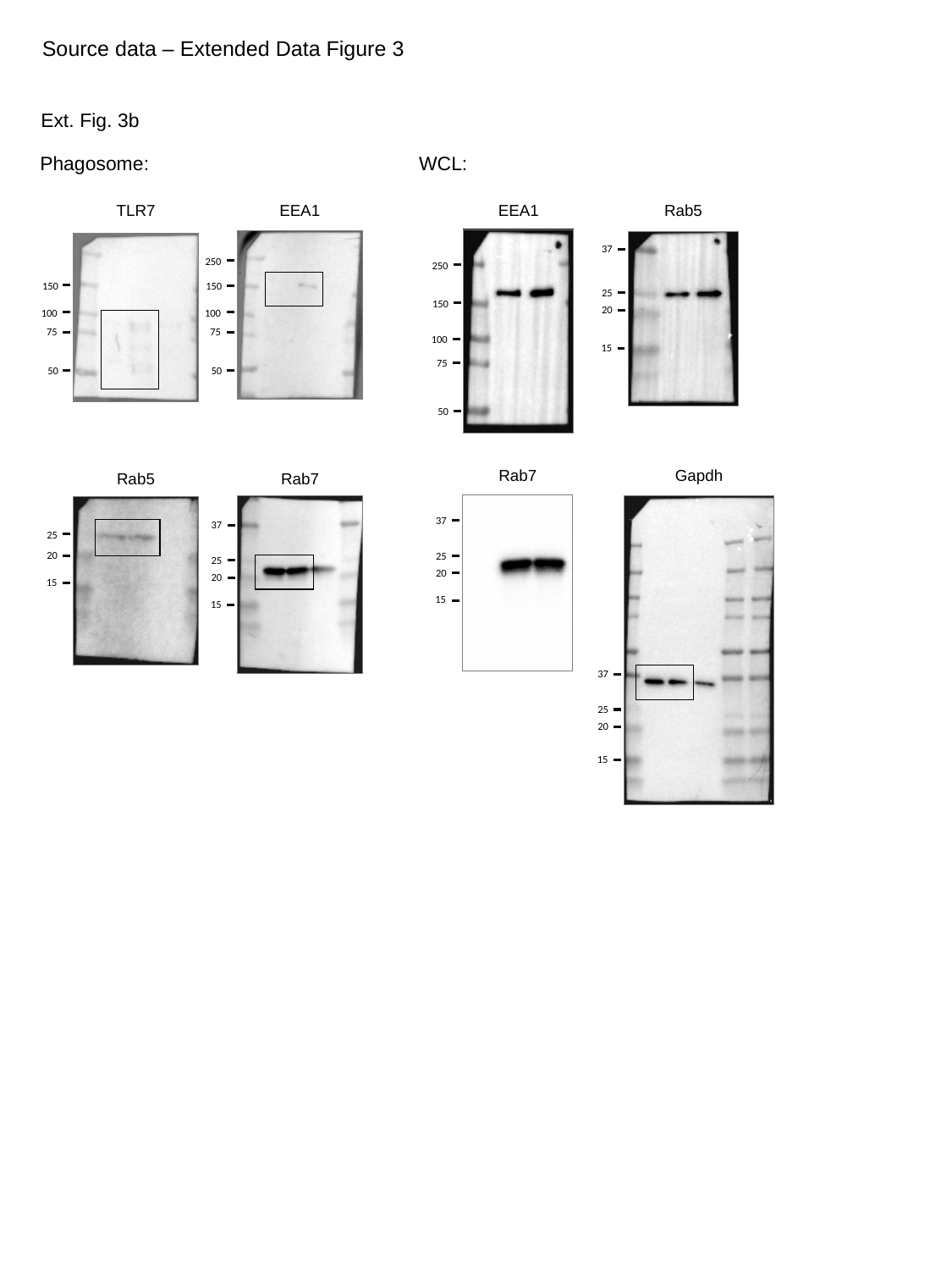

Source data – Extended Data Figure 3
Ext. Fig. 3b
Phagosome:
WCL:
TLR7
EEA1
EEA1
Rab5
37
250
250
150
150
25
150
20
100
100
75
75
100
15
75
50
50
50
Rab7
Gapdh
Rab5
Rab7
37
37
25
20
25
25
20
20
15
15
15
37
25
20
15

### Slide 6
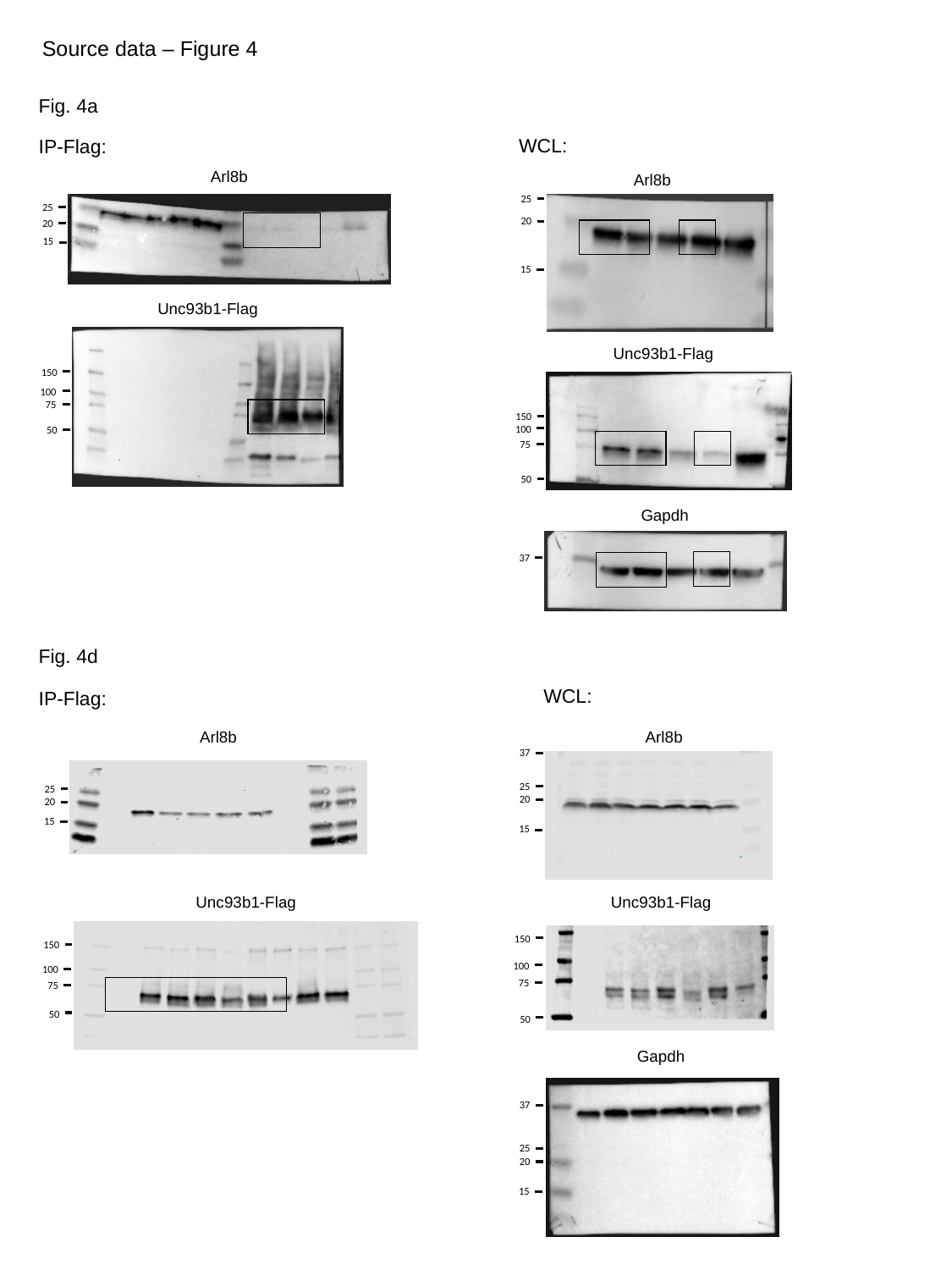

Source data – Figure 4
Fig. 4a
WCL:
IP-Flag:
Arl8b
Arl8b
25
25
20
20
15
15
Unc93b1-Flag
Unc93b1-Flag
150
100
75
150
100
50
75
50
Gapdh
37
Fig. 4d
WCL:
IP-Flag:
Arl8b
Arl8b
37
25
25
20
20
15
15
Unc93b1-Flag
Unc93b1-Flag
150
150
100
100
75
75
50
50
Gapdh
37
25
20
15

### Slide 7
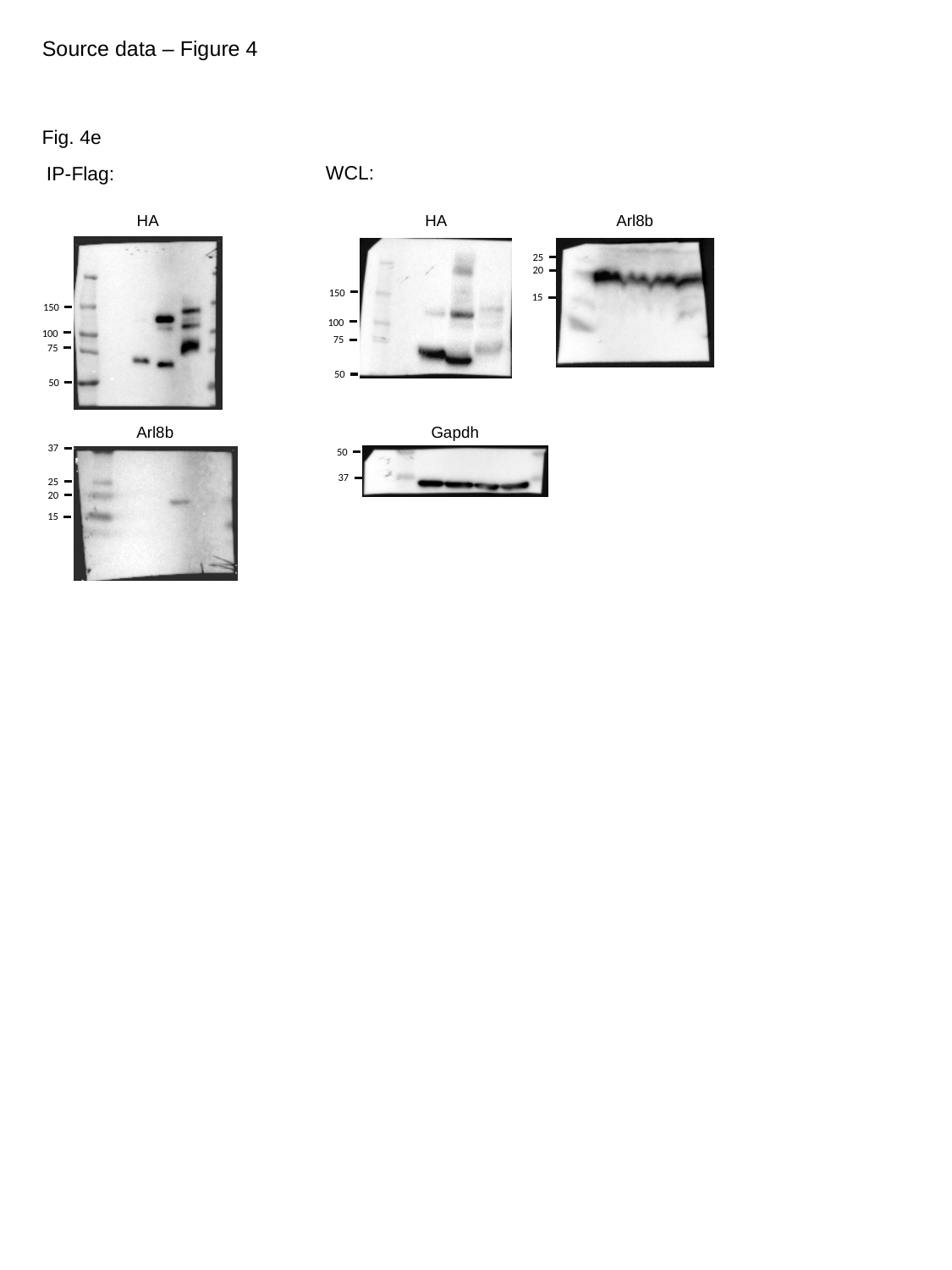

Source data – Figure 4
Fig. 4e
WCL:
IP-Flag:
HA
HA
Arl8b
25
20
150
15
150
100
100
75
75
50
50
Arl8b
Gapdh
37
50
37
25
20
15

### Slide 8
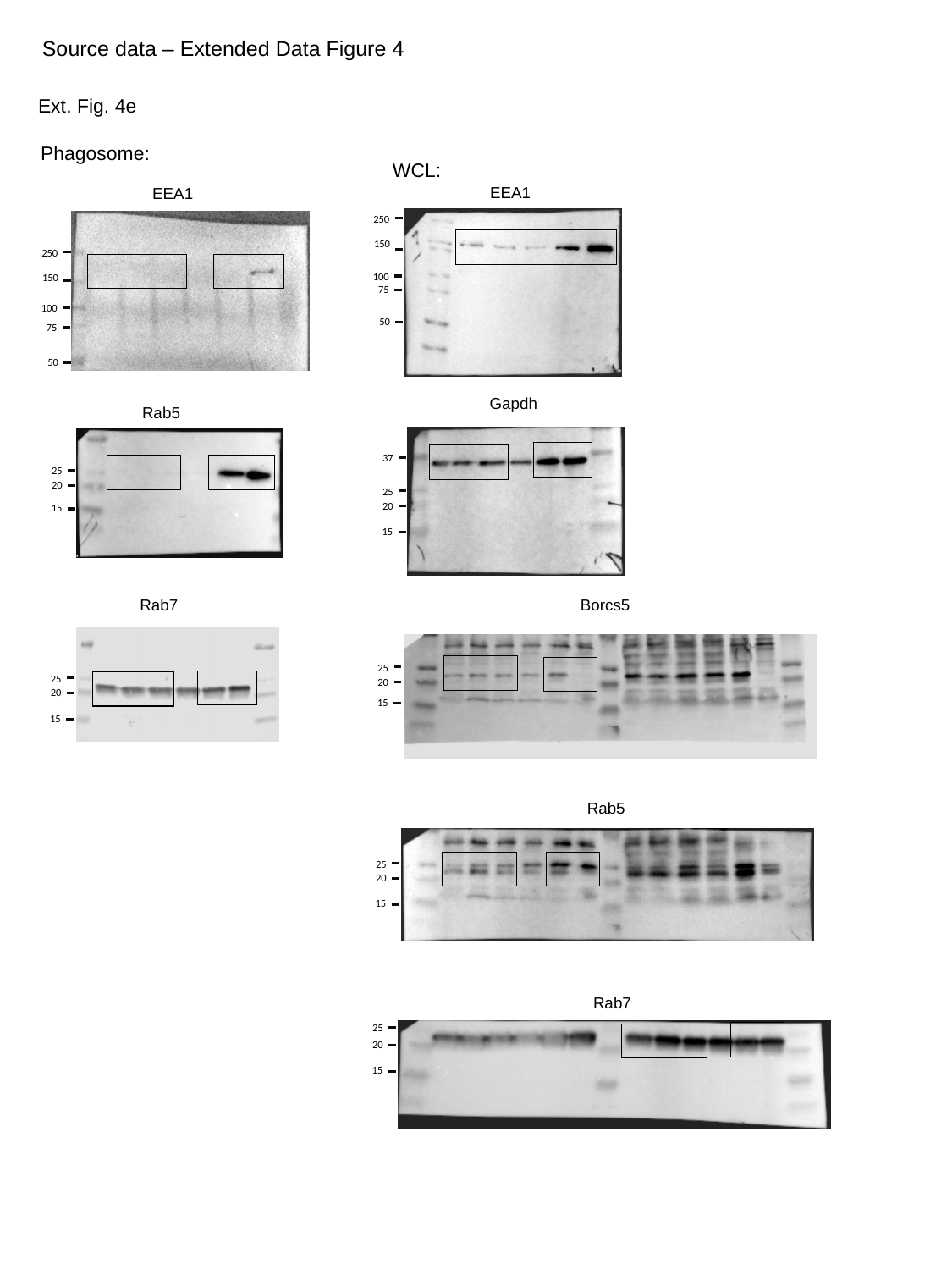

Source data – Extended Data Figure 4
Ext. Fig. 4e
Phagosome:
WCL:
EEA1
EEA1
250
150
250
100
150
75
100
50
75
50
Gapdh
Rab5
37
25
20
25
20
15
15
Borcs5
Rab7
25
25
20
20
15
15
Rab5
25
20
15
Rab7
25
20
15

### Slide 9
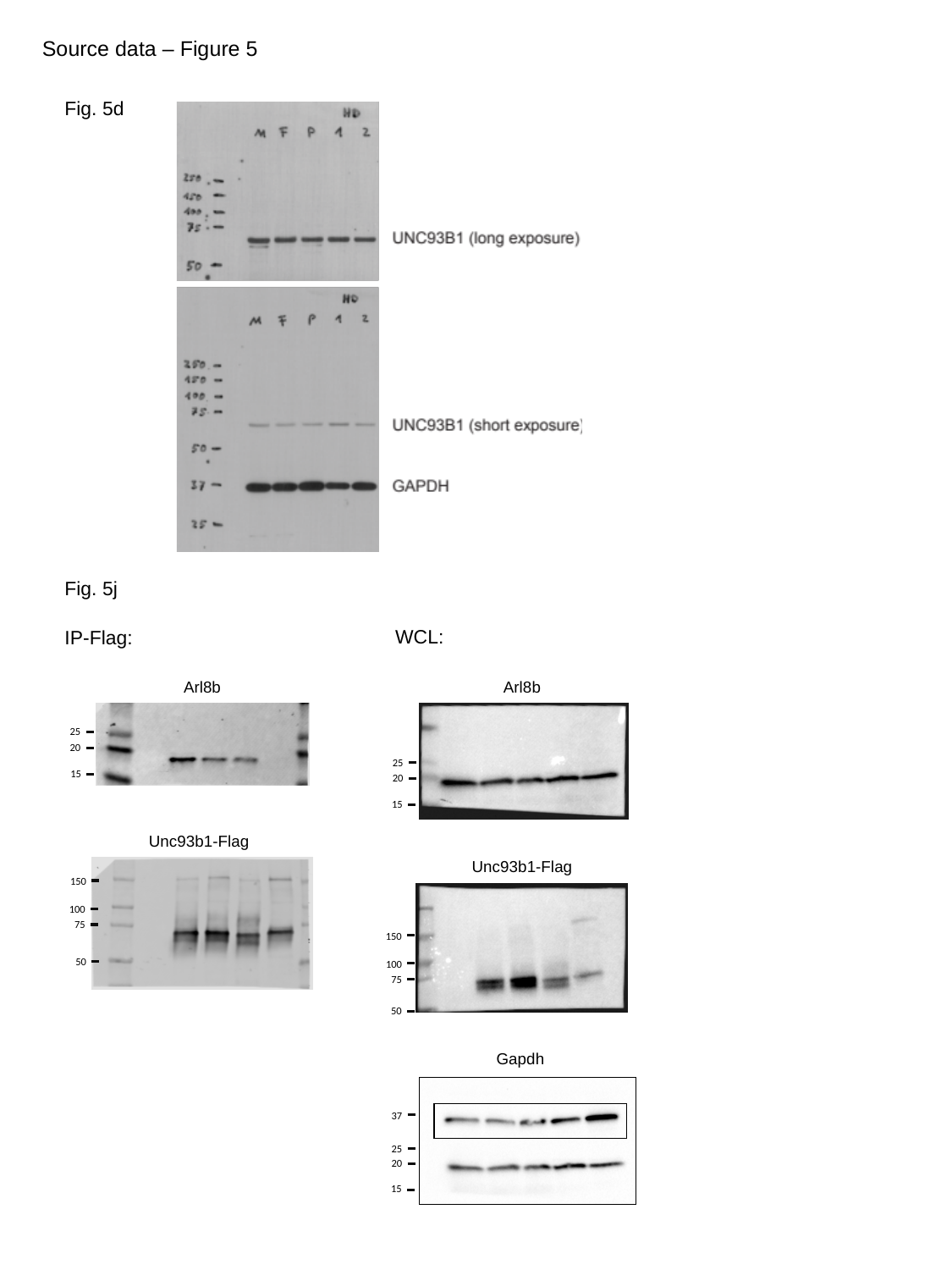

Source data – Figure 5
Fig. 5d
Fig. 5j
WCL:
IP-Flag:
Arl8b
Arl8b
25
20
25
15
20
15
Unc93b1-Flag
Unc93b1-Flag
150
100
75
150
50
100
75
50
Gapdh
37
25
20
15

### Slide 10
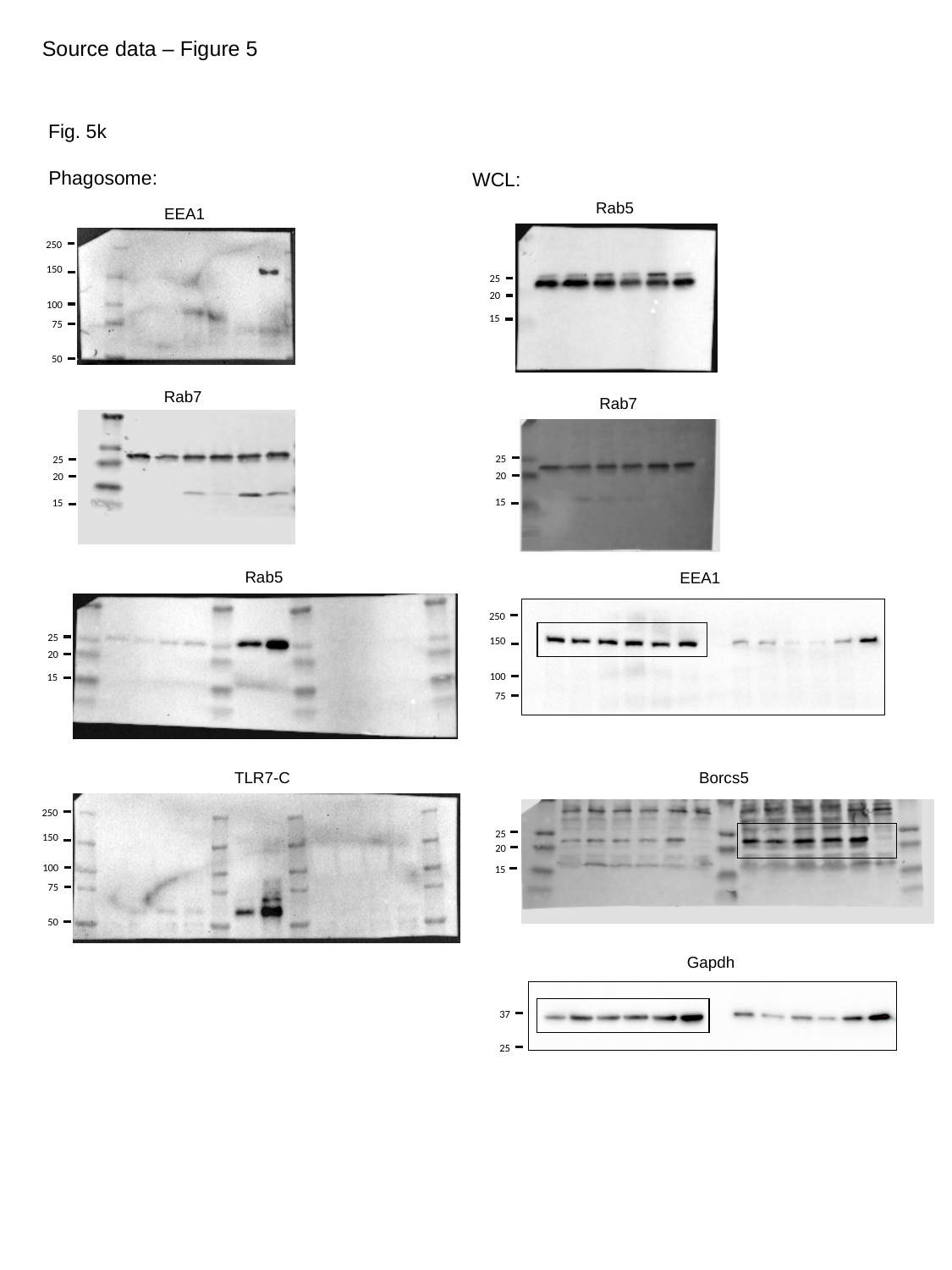

Source data – Figure 5
Fig. 5k
Phagosome:
WCL:
Rab5
EEA1
250
150
25
20
100
15
75
50
Rab7
Rab7
25
25
20
20
15
15
Rab5
EEA1
250
25
150
20
100
15
75
Borcs5
TLR7-C
250
25
150
20
100
15
75
50
Gapdh
37
25
